# Pex3 promotes formation of peroxisome-peroxisome and peroxisome-lipid droplet contact sites

**DOI:** 10.1101/2024.12.10.627744

**Authors:** Lucía Amado, Rico Franzkoch, Louis Percifull, Vico Flatemersch, Eleni Joana Brüggermann, Olympia Ekaterini Psathaki, Maya Schuldiner, Maria Bohnert, Margret Bülow, Ayelén González Montoro

## Abstract

Peroxisomes are ubiquitous organelles that mediate central metabolic functions, such as fatty acid β-oxidation, as well as diverse tissue- and organism-specific processes. Membrane contact sites, regions of close apposition with other organelles for direct communication, are central to several aspects of their life cycle. Pex3 is a conserved multifunctional peroxisomal transmembrane protein involved in the insertion of peroxisomal membrane proteins, in pexophagy, and in the formation of membrane contact sites. Here, we show that elevated Pex3 levels in *Saccharomyces cerevisiae* induce the formation of peroxisome clusters surrounded by lipid droplets, mediated by peroxisome-peroxisome and peroxisome-lipid droplet contact sites. This clustering occurs independently of Pex3 partners in other processes, Pex19, Inp1, and Atg36. The cytosolic domain of Pex3 binds peroxisomes, suggesting a direct role in homotypic contact site formation. Lipid droplet-peroxisome contact sites require the lipid droplet-localized triacylglycerol lipase Tgl4, which is enriched along with other lipases at this interface. Pex3 overexpression in *Drosophila melanogast*er similarly alters peroxisome and lipid droplet morphology and promotes contact site formation. Together, our results offer novel molecular insights into homotypic peroxisome contact sites and peroxisome-lipid droplet contact sites across species.

## INTRODUCTION

Peroxisomes are found in most eukaryotic cells and are the place of important metabolic reactions, including the oxidation of fatty acids, the synthesis of ether lipids, and the detoxification of hydrogen peroxide and glyoxylate. While some peroxisomal functions are tissue or organism-specific, others are highly conserved, e.g. the oxidation of fatty acids, a virtually ubiquitous process (Wanders et al., 2023). Impaired peroxisome biogenesis and defects in peroxisomal metabolic pathways result in severe human diseases collectively known as peroxisomal disorders (Wanders et al., 2023).

Membrane contact sites are structurally defined regions of close organelle apposition without membrane fusion (Scorrano et al., 2019; Eisenberg-Bord et al., 2016). A central function of contact sites is the exchange of material among the compartments, including the transport of luminal material and of membrane lipids. Furthermore, the physical attachment of the organelle membranes can affect organelle fusion, fission, and positioning (Prinz et al., 2020). Membrane contact sites exist between virtually all pairs of organelles (Valm et al., 2017, Shai et al., 2018, Kakimoto et al., 2018), and coordinate multi-organelle processes (Prinz et al., 2020; Voeltz et al., 2024). Contact sites between peroxisomes and other organelles play important roles in different aspects of the peroxisome life cycle (Shai et al., 2016). For example, in the yeast *Saccharomyces cerevisiae*, peroxisome contact sites with the endoplasmic reticulum affect peroxisome proliferation (David et al., 2013; Yan et al., 2008), and the formation of peroxisome contact sites with the cell periphery determines the inheritance of these organelles among mother and daughter cells (Knoblach et al., 2013a).

Pex3 is a peroxisomal membrane protein with multiple functions and interactors. It is involved in the targeting of peroxisomal membrane proteins, by acting as a docking factor for Pex19, a cytosolic receptor for peroxisomal membrane protein precursors. Another function of Pex3 is the recruitment of the pexophagy receptor Atg36. Both of these functions are conserved in the human Pex3 protein (Burnett et al., 2015; Fang et al., 2004; Motley et al., 2012; Sato et al., 2008, 2010; Schmidt et al., 2012; Yamashita et al., 2014). Pex3 has additional roles in contact site formation. It acts as a membrane anchor for Inp1, a tethering protein involved in the retention of peroxisomes in the mother cell during cell division, through its interaction with the cell cortex (Hulmes et al., 2020; Knoblach et al., 2013a). In the yeast *Hansenula polymorpha* extensive vacuole-peroxisome contact sites are formed when cells are shifted from glucose containing media to methanol containing media, a transition that requires peroxisomal expansion. Pex3 is enriched at these contact sites, and overexpression of the protein results in contact site expansion during growth in glucose, suggesting that Pex3 might be a tether (Wu et al., 2019).

This prompted us to address the effects of Pex3 overexpression in *Saccharomyces cerevisiae*. We find that cells that overexpress Pex3 contain peroxisome clusters surrounded by lipid droplets, which are formed by peroxisome-peroxisome and peroxisome-lipid droplet contact sites. We further show that these contact sites are independent of all the known Pex3 interactors. Instead, efficient formation of peroxisome-lipid droplet contact sites requires the lipid droplet-localized TAG-lipase Tgl4 but not other lipid droplet lipases, even though several of them are enriched in the interface. Interestingly, similar effects of Pex3 overexpression on peroxisome morphology and extended contact with the lipid droplets were also observed in *Drosophila melanogaster*, showing that these aspects of Pex3 function are conserved to metazoa. Altogether our findings expand our understanding of peroxisomal contact sites, and of the multi-function protein Pex3.

## RESULTS

### Overexpression of Pex3 causes a change in the morphology of peroxisomes and lipid droplets

In *Hansenula polymorpha*, Pex3 was observed to be enriched in membrane contact sites between peroxisomes and the vacuole, and overexpression of this protein results in expansion of these contact sites (Wu et al., 2019). Thus, we decided to test the phenotype of Pex3 overexpression in *Saccharomyces cerevisiae.* Overexpression from the strong constitutive *TEF1* promoter resulted in a morphological change in peroxisomes marked by mCherry directed to the peroxisomal lumen by a PTS1 signal consisting of a serine-lysine-leucine sequence (mCh-SKL). While control cells show on average 5.5 mCh-SKL positive structures, cells overexpressing Pex3 contain mainly one structure (Figure 1A and B). This structure was found in close proximity to the vacuole in 55% of cells. We confirmed these observations by using other peroxisomal markes, Pex3 itself and Pex14, obtaining similar results (Supplemental Figure 1A and B).

**Figure 1:**
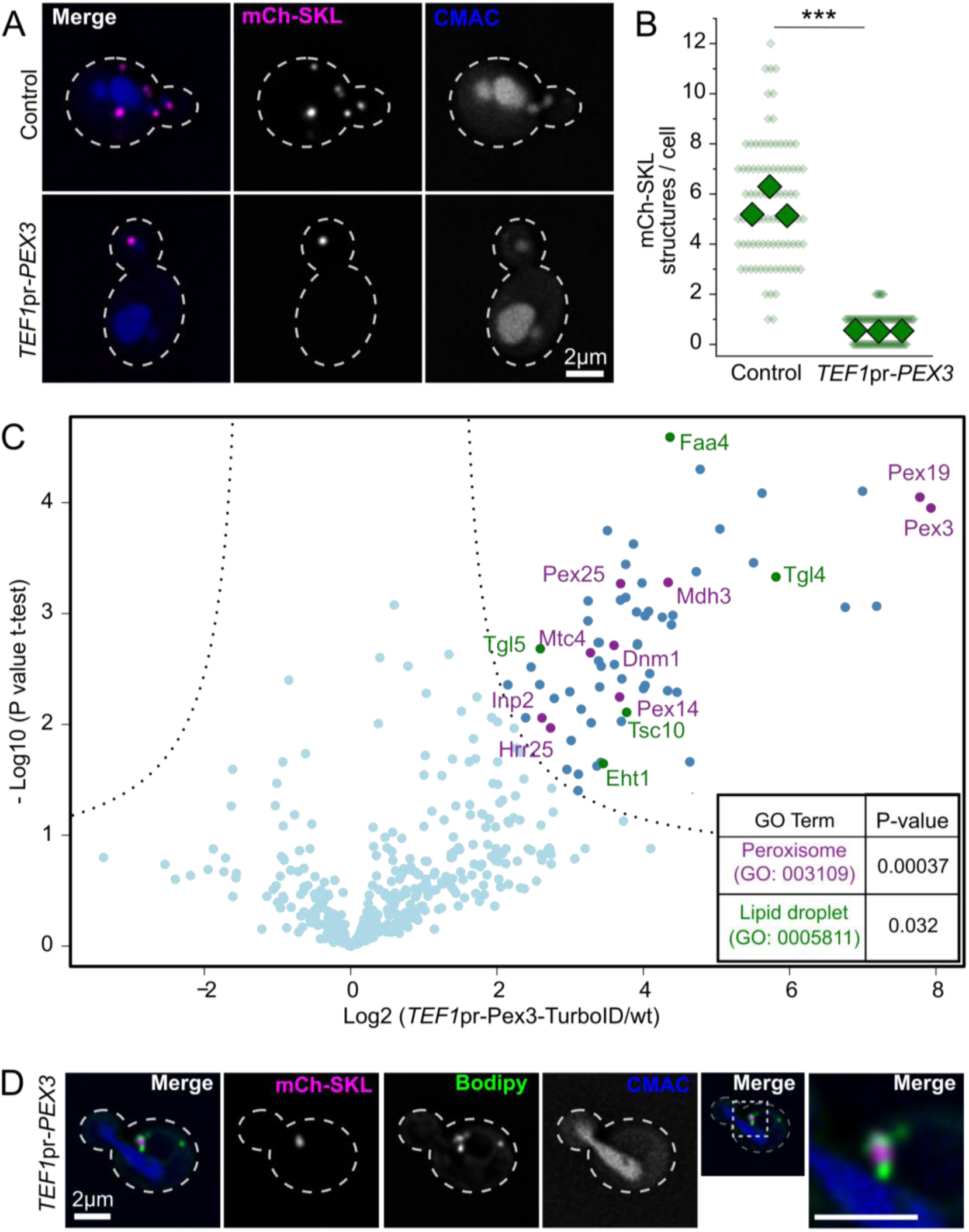
Overexpression of Pex3 causes the formation of a single peroxisomal structure surrounded by lipid droplets. A-B) Overexpression of Pex3 produces collapse of all peroxisomal signal into one structure. Panel A shows representative fluorescence microscopy images of a strain expressing mCherry-SKL construct to visualize the lumen of the peroxisomes, either with Pex3 at endogenous levels (Control) or overexpressed (*TEF1pr-PEX3*) and the vacuolar lumen stained with CMAC. Cell outlines are shown as white dashed lines. Scale bar: 2 μm. Panel B shows the quantification of the amount of peroxisomal structures per cell. Three independent experiments were performed and 30 cells were analyzed for each experiment and condition. Small diamonds correspond to individual cells, bigger diamonds correspond to the average of independent experiments. The different strains were compared using an unpaired two-tailed Student’s t-test. *** P < 0.001. C) Turbo ID of overexpressed Pex3-TurboID enriches peroxisomal and lipid droplets proteins. Volcano plot showing relative protein intensity in a pull-down of biotinylated proteins between a strain overexpressing Pex3 tagged c-terminally with the TurboID protein (*TEF1pr-PEX3-TID*) and a wild type control strain (wt). Peroxisomal proteins are marked in magenta and lipid droplet proteins in green. GO term enrichment analysis of the group of proteins significantly enriched in the Pex3-TID pull-down showed an enrichment of the GO Terms “peroxisomes” and “lipid droplets”. D) The formed peroxisomal structure is surrounded by lipid droplets. Representative fluorescence microscopy image of a strain expressing mCherry-SKL construct to visualize the lumen of the peroxisomes, with Pex3 overexpressed (*TEF1pr-Pex3*), the vacuolar lumen is stained with CMAC and lipid droplets are stained with Bodipy. Zoomed in region shows a peroxisomal structure surrounded by lipid droplets. Cell outlines are shown as white dashed lines. Scale bar: 2 μm.

To characterize this structure we sought to describe its molecular microenvironment by proximity biotinylation (Fernández-Suárez et al., 2008), by tagging Pex3 with TurboID (Branon et al., 2018) at its C-terminus, which faces the cytosol. Cells expressing Pex3-TurboID under the *TEF1* promoter were incubated with biotin for 3 hours and the biotinylated proteins were isolated by affinity chromatography using a streptavidin matrix. The bait protein Pex3 was among the most highly enriched proteins, as was its known interactor Pex19 (Figure 1C). A Gene Ontology (GO) Term enrichment analysis showed the GO term “Peroxisome” as the most represented annotation among our enriched proteins, as expected. Interestingly, the GO term “Lipid droplet” was also significantly enriched (P value = 0.032, Figure 1C).

This prompted us to address the subcellular localization of lipid droplets when Pex3 is overexpressed. This analysis revealed that under these conditions, lipid droplets are strongly recruited to the peroxisomal structure, with 95% of these structures being in close proximity to or surrounded by lipid droplets (Figure 1D). We conclude that overexpression of Pex3 causes a change in the morphology of peroxisomes and lipid droplets, resulting in the observation of a single peroxisomal structure per cell, which is in close proximity to lipid droplets.

### Pex3 overexpression induces a cluster of peroxisomes surrounded by lipid droplets, which includes peroxisome-peroxisome and peroxisome-lipid droplet contact sites

To understand the characteristics of the structure formed by lipid droplets and peroxisomes upon Pex3 overexpression, we sought to enlarge the structure to gain spatial resolution. This was achieved by deleting *PEX11*, which results in enlarged peroxisomes (Erdmann & Blobel, 1995; Yifrach et al., 2022). Additionally, we cultured the cells in the presence of oleate, which resulted in enlarged lipid droplets. These conditions allowed sufficient spatial resolution to distinguish the peroxisomal matrix from the membrane, as peroxisomal membrane proteins (Pex13 and Pex3) were observed to surround BFP directed to the peroxisomal lumen by a C-terminal PTS1 signal (BFP-SKL). Under these conditions, BFP-SKL was observed as a cluster with multiple maxima. The peroxisomal membrane proteins showed peaks between the individual maxima of the BFP-SKL signal (Figure 2A and B). This indicates that the peroxisomal structure formed upon Pex3 overexpression corresponds to a cluster of multiple peroxisomes. The lipid droplets marked by Erg6-2xmKate2 were observed to surround the cluster of peroxisomes.

**Figure 2:**
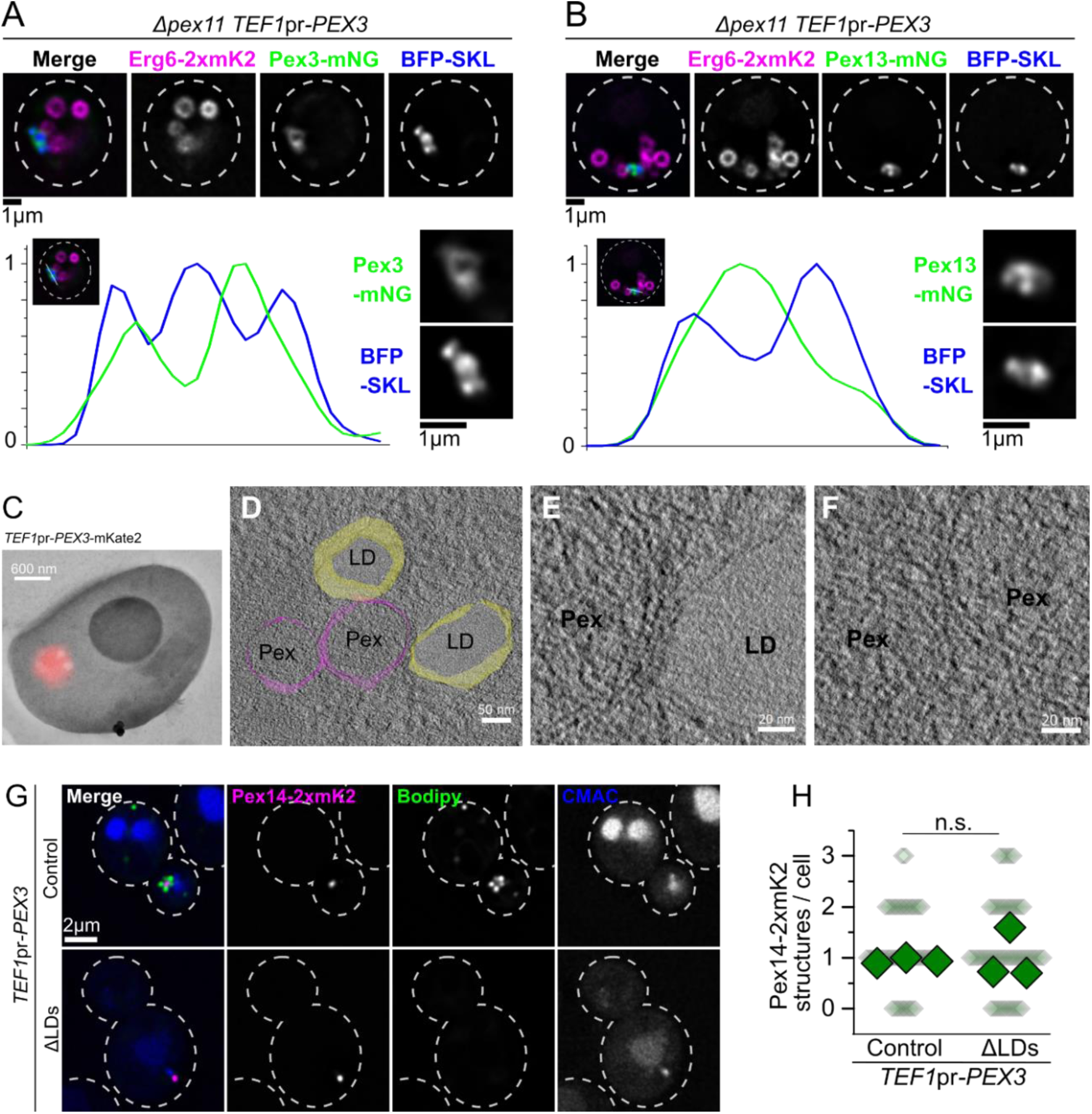
Overexpression of Pex3 induces an accumulation of peroxisomes surrounded by LDs that contains Pex-Pex and Pex-LD contact sites. A-B) Enlarged peroxisomes and LDs reveal that the structures contain several maxima for peroxisome lumen signal, with peroxisomal membrane between them. Representative pictures of a strain with overexpressed Pex3 (TEF1pr-PEX3), expressing the BFP-SKL construct to visualize the lumen of the peroxisomes, Erg6-2xmKate2 marking the lipid droplet monolayer, and either Pex3 (A) or Pex13 (B) tagged with mNeonGreen as markers of the peroxisomal membrane. To produce enlarged peroxisomes and lipid droplets, the cells contain a deletion of PEX11, and were grown with oleate as the sole carbon source for 20hs. Cell outlines are shown as white dashed lines. Scale bars: 1 μm. Each graph shows the signal of BFP-SKL and the corresponding peroxisomal membrane protein over a line across the structure, depicted in the merged image. C-F) On-section CLEM tomography confirms that the structure involves Pex-Pex and Pex-LD contact sites. Panel C shows a representative image of on-section CLEM done on a strain with overexpressed Pex3 (TEF1pr-PEX3) and tagged with 2xmKate2. Scale bar: 600 nm. Panel D shows a tomography image of the same section overlayed with the 3D model recreated from the images, showing peroxisomes in magenta and lipid droplets in yellow. Scale bar: 50 nm. Panels E and F show zoomed in regions of the tomogram as examples of the peroxisome-lipid droplet (E) and peroxisome-peroxisome (F) contact sites. Scale bars: 20 nm. G-H) LDs are not necessary for the formation of the cluster of peroxisomes. Panel G shows representative pictures of strains with overexpressed Pex3 (TEF1pr-PEX3) in control cells and in strains that cannot produce lipid droplets (ΔLDs), expressing Pex14-2xmKate2 to visualize the peroxisomes, lipid droplets were stained with Bodipy and the vacuolar lumen was stained with CMAC. Cell outlines are shown as white dashed lines. Scale bar: 2 μm. Panel H shows the quantification of the amount of peroxisomal structures per cell. Three independent experiments were performed and 30 cells were analyzed for each experiment and condition. Small diamonds correspond to individual cells, bigger diamonds correspond to the average of independent experiments. The different strains were compared using an unpaired two-tailed Student’s t-test. n.s., not significant.

Given the close and specific proximity observed between these organelles upon overexpression of Pex3, we reasoned that such an organelle cluster would likely be formed by peroxisome-peroxisome and peroxisome-lipid droplet contact sites. To test this, we performed on-section CLEM tomography of cells overexpressing Pex3-mKate2, using the mKate2 signal to locate the structures. This approach revealed clusters of peroxisomes surrounded by lipid droplets that included peroxisome-peroxisome and peroxisome-lipid droplet contact sites (Figure 2C - D). Figure 2C shows a on-section correlative fluorescence microscopy and TEM image from a cell overexpressing Pex3-mKate2, used to identify the relevant region for TEM tomography. Figure 2D shows imaging of this region by TEM tomography, overlayed with a 3D model reconstructing the observed organelles. Figures 2E and F show example regions of the tomogram, containing peroxisome-lipid droplet and peroxisome-peroxisome contact sites.

We asked if the formation of peroxisome-peroxisome contact sites and peroxisome-lipid droplet contact sites occurred independently, or whether the clustering of the two types of organelles was interlinked. To test this, we overexpressed Pex3 in a strain that is devoid of lipid droplets because it lacks the synthases for triglycerides and sterol esters, namely Dga1, Lro1, Are1 and Are2 (from here on termed ΔLDs) (Sandager et al., 2002). Overexpression of Pex3 caused the accumulation of peroxisomal signal into a single structure irrespective of the absence of lipid droplets, showing that lipid droplets are not required for the formation of the peroxisome-peroxisome contact sites (Figure 2G and H).

### The cytosolic domain of Pex3 interacts with peroxisomes

Pex3 is anchored to the peroxisomal membrane by a single transmembrane domain at its N-terminus, which is sufficient to cause targeting to the peroxisomes, and contains a globular cytosolic C-terminal domain (Höhfeld et al., 1991; Soukupova et al., 1999; Kammerer et al., 1998; Schmidt et al., 2010) (Figure 3A). To address a possible direct involvement of Pex3 in contact site formation, we expressed the cytosolic domain (CD – amino acids 40-441) fused to GFP at its N-terminus and lacking the transmembrane region (Figure 3A). We observed that this construct localized at peroxisomes, marked by Pex14-HaloTag (Figure 3B). The construct did not decorate lipid droplets marked by Erg6-2xmKate2, and was only observed enriched in these structures when peroxisomal signal was also present (Figure 3B, see zoomed in organelles). Consistently, calculation of the Mandeŕs coefficients M1 and M2 for GFP-Pex3(CD) with Pex14-HaloTag showed high values, while the coefficients of overlap with the Erg6-2xmKate2 signal were much lower and comparable to the ones observed between Pex14-HaloTag and Erg6-2xmKate2 (Figure 3C).

**Figure 3:**
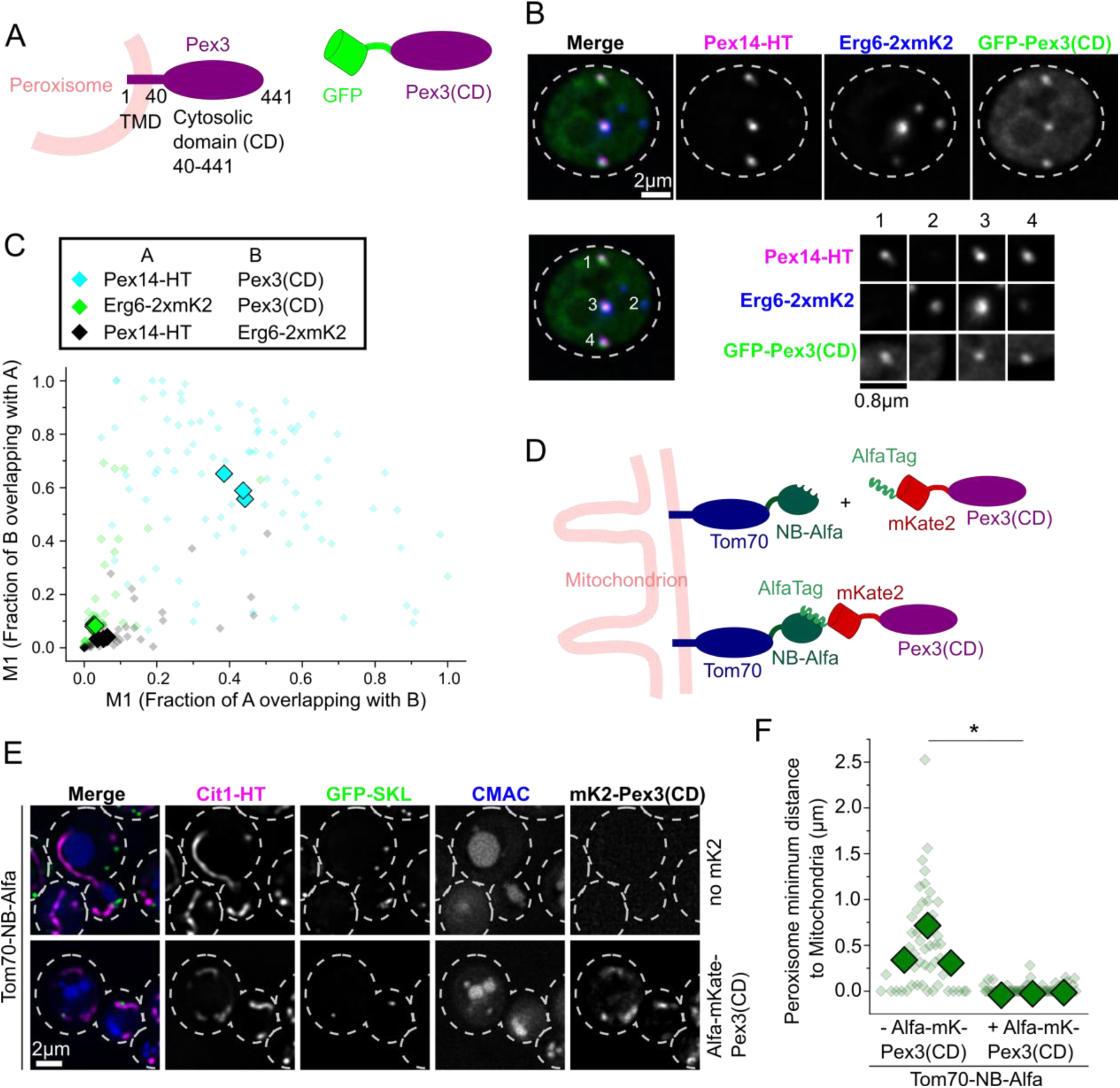
The cytosolic domain of Pex3 binds peroxisomes and is able to tether them to another organelle. A-C) The cytosolic domain of Pex3 binds to peroxisomes. Panel A shows a diagram of the GFP-Pex3(CD) construct. This construct contains the cytosolic domain of Pex3 (aa40-441) fused to GFP tag at the N-terminus instead of the transmembrane domain as in the full length Pex3. Panel B shows representative pictures of the colocalization experiment of GFP-Pex3(CD) with peroxisomes (Pex14-HaloTag) and lipid droplets (Erg6-2xmKate2). Cell outlines are shown as white dashed lines. Scale bar: 2 μm and 0.8 μm. Panel C shows the co-localization analysis of the experiment in B using Mandeŕs coefficients M1 and M2 for the overlap of: Pex14-HaloTag and GFP-Pex3(CD) (cyan diamonds), Erg6-2xmKate2 and GFP-Pex3(CD) (green diamonds) or Pex14-HaloTag and Erg6-2xmKate2 (black diamonds). Three independent experiments were performed and 30 cells were analyzed for each experiment. Each small diamond represents a single cell, and the bigger ones represent the average of each of three independent experiments. D) Diagram of the strategy used to recruit the cytosolic domain of Pex3 to mitochondria artificially. The outer mitochondrial membrane protein Tom70 was tagged in the C-terminus with a Nanobody that recognizes the AlfaTag (Tom70-NB-Alfa). The cytosolic domain of Pex3 was tagged with an AlfaTag and an mKate2 fluorescent protein in the N-terminus (AlfaTag-mKate2-Pex3(CD) construct). This causes the recruitment of AlfaTag-mKate2-Pex3(CD) to mitochondria. E-F) Targeting the cytosolic domain of Pex3 to mitochondria tethers peroxisomes to this organelle. Panel E shows representative images of the localization of peroxisomes (GFP-SKL) and mitochondria (Cit1-HaloTag) in the presence or absence of AlfaTag-Pex3(CD), with Tom70 fused to Alfa Nanobody in the background. Cell outlines are shown as white dashed lines. Scale bar: 2 μm. Panel F shows the measurements of distances between peroxisomes (GFP-SKL) and mitochondria (Cit1-HaloTag) in the presence or absence of the AlfaTag-Pex3(CD) construct as described. Three independent experiments were performed and 30 cells were analyzed for each experiment. Each small diamond represents a single peroxisome, and the bigger ones represent the average of each of three independent experiments. Strains were compared using the means of each experiment, with an unpaired, two-tailed Student’s t-test. * P < 0.05.

To test if the interaction of the cytosolic domain of Pex3 with peroxisomes is strong enough to enable organelle tethering, we artificially directed this domain to mitochondria. This was achieved by tagging it with the fluorescent protein mKate2 and an N-terminal Alfa tag (Götzke et al., 2019). In addition, the mitochondrial outer membrane receptor Tom70 was tagged with a nanobody that recognizes the Alfa tag (Götzke et al., 2019), so it should direct the Pex3(CD) to mitochondria (Figure 3D). Indeed Alfa-mKate2-Pex3(CD) decorates the mitochondrial network under these conditions, as confirmed by co-localization with the mitochondrial marker Cit1-HaloTag (Figure 3E). This caused peroxisomes to re-localize and attach to the surface of mitochondria (Figure 3E and F), confirming that the interaction of Pex3(CD) with peroxisomes is strong enough to induce organelle tethering. Based on these results we propose that the homotypic peroxisomal contact sites are formed by Pex3 being anchored to the peroxisomal membrane via its transmembrane domain, and additionally interacting with other peroxisomes through its cytosolic domain.

### Known interactors of Pex3 are not involved in forming the peroxisome-lipid droplet cluster

Pex3 is a multifunctional protein involved in different processes related to the life-cycle of peroxisomes, including the targeting of peroxisomal membrane proteins, pexophagy, and targeting of peroxisomes to the cortex, which strongly influences peroxisome inheritance (Burnett et al., 2015; Fang et al., 2004; Hulmes et al., 2020; Knoblach et al., 2013; Motley et al., 2012; Sato et al., 2008, 2010; Yamashita et al., 2014). These functions are mediated by the direct interaction of Pex3 with different binding partners (Figure 4A). Targeting of peroxisomal membrane proteins to the peroxisomal membrane involves its interaction with the cytosolic receptor Pex19 (Fang et al., 2004; Sato et al., 2010) while its role in autophagy is mediated by its interaction with the pexophagy receptor Atg36 (Motley et al., 2012). Finally, the tethering of peroxisomes to the cell cortex is mediated by Pex3 interacting with Inp1 (Hulmes et al., 2020; Knoblach et al., 2013) (Figure 4A). Next, we tested if any of the known interactors of Pex3 is involved in the formation of the peroxisome-lipid droplet cluster. To do this, we overexpressed Pex3 in strains lacking either *ATG36* or *INP1.* The representative microscopy images and quantifications shown in Figures 4B - G show that the phenotype caused by Pex3 overexpression in these backgrounds does not differ from the control cells, indicating that neither Inp1 nor Atg36 are required for the formation of this structure.

**Figure 4:**
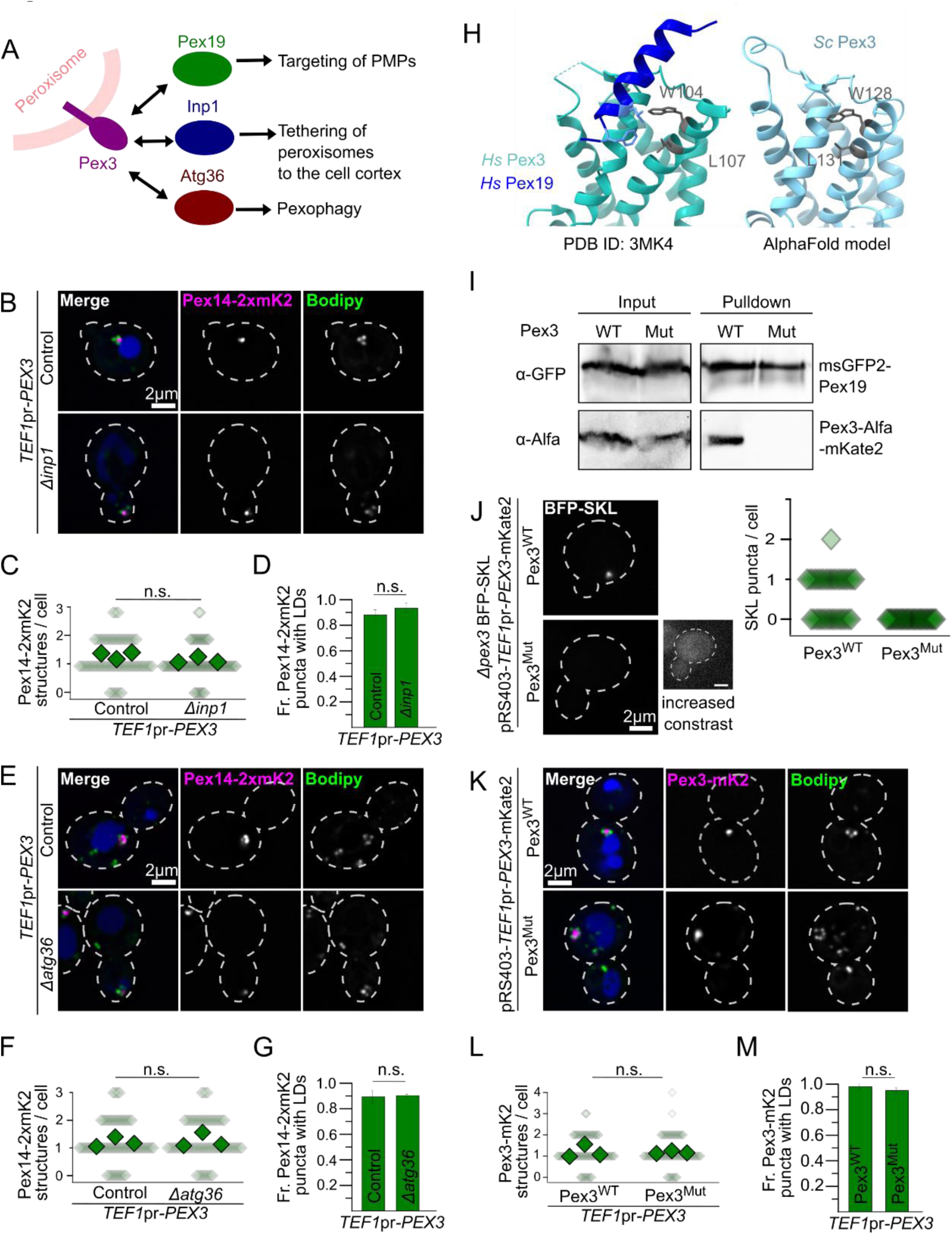
Formation of the peroxisome-lipid droplet cluster is independent of known interactors of Pex3. A) Diagram of Pex3 known interactors and their functions. B, E) Representative images of strains overexpressing Pex3 (TEF1pr-PEX3) in control cells and strains lacking Inp1 (B) or Atg36 (E). All strains express Pex14 fused to 2xmKate2 to visualize the peroxisomes, lipid droplets were stained with Bodipy and the vacuolar lumen was stained with CMAC. Cell outlines are shown as white dashed lines. Scale bars: 2 μm C, F) Quantification of the number of peroxisomal structures per cell in the microscopy experiments described before. Small diamonds correspond to individual cells, bigger diamonds correspond to the average of independent experiments. Three independent experiments were performed and 30 cells were analyzed for each experiment and condition. The different strains were compared using an unpaired two-tailed Student’s t-test. n.s., not significant. D, G) Quantification of the fraction of peroxisomal structures with accumulations of lipid droplets next to them in the microscopy experiments described above. Three independent experiments were performed and 30 cells were analyzed for each experiment and condition. The different strains were compared using an unpaired two-tailed Student’s t-test. n.s., not significant.H) The amino acids involved in hsPex3 interaction with hsPex19 are conserved in yeast. To the left, the structure obtained for Homo sapiens Pex3 (cyan) interacting with a peptide of hsPex19 (dark blue) (Sato et al., 2010). Amino acids W104 and L107 of hsPex3 are involved in the interaction with hsPex19. To the right, the structure predicted by AlphaFold for Saccharomyces cerevisiae Pex3 (light blue) shows that it contains a structurally conserved W and L in the same positions (W128 and L131 in scPex3). I) Pex3(W128K, L131K) cannot interact with Pex19. Affinity purification of msGFP2-Pex19 co-purifies Pex3-mKate2-AlfaTag but not Pex3(W128K, L131K)-mKate2-AlfaTag. J) Pex3(W128K, L131K) does not support BFP-SKL import into peroxisomes. In a strain that expresses BFP-SKL, endogenous Pex3 was deleted and either Pex3wt or Pex3(W128K, L128K) were re-introduced in a plasmid. Quantification of the number of BFP-SKL puncta per cell is shown to the right. K-M) Overexpression of Pex3(W128K, L131K), which cannot interact with Pex19, still causes aggregation of peroxisomes and recruitment of lipid droplets. Panel K shows representative images of strains overexpressing Pex3 from a plasmid (TEF1pr-PEX3-mKate2) containing either Pex3 wt or Pex3 mutant. Lipid droplets were stained with Bodipy and the vacuolar lumen was stained with CMAC. The strain contains Pex3 wt in the background, expressed from its genomic locus, in order to have normal peroxisomes. Cell outlines are shown as white dashed lines. Scale bars: 2 μm. Panel L shows the quantification of the number of peroxisomal structures per cell in the microscopy experiments described before. Small diamonds correspond to individual cells, bigger diamonds correspond to the average of independent experiments. Panel M shows the quantification of the proportion of peroxisomal structures with accumulations of lipid droplets next to them in the microscopy experiments described above. Three independent experiments were performed and 30 cells were analyzed for each experiment and condition. The different strains were compared using an unpaired two-tailed Student’s t-test. n.s., not significant.

To test the requirement of *PEX19*, we could not delete this gene, as this results in the absence of functional peroxisomes (Hettema et al., 2000). We thus used an alternative strategy, by addressing whether overexpression of a mutant version of Pex3 that does not interact with Pex19 still promotes the formation of the peroxisome-lipid droplet cluster. The crystal structure of human Pex3 in complex with a fragment of human Pex19 has previously been solved. It was found that the interaction with Pex19 is mediated by a region centered around HsPex3-Trp104, which also includes Leu107 (Sato et al., 2008; F. Schmidt et al., 2010). Alignment of the AlphaFold-generated structure prediction of ScPex3 with the structure of HsPex3 indicated that this region is highly conserved and that the equivalent residues in ScPex3 are Trp128 and Leu131 (Figure 4H). We thus introduced the mutations Trp128Lys and Leu131Lys in ScPex3 (Pex3^Mut^) and tested the ability of this mutant to interact with Pex19, support peroxisome biogenesis, and form the peroxisome-lipid droplet structures upon overexpression. Unlike wtPex3, Pex3^Mut^ was not co-purified with msGFP2-Pex19, indicating that the mutations disrupt this interaction (Figure 4I). The Pex19-Pex3 interaction has been reported to be required to import PTS1 containing peroxisomal matrix proteins (Fang et al., 2004). Consistently, we observed that expression of Pex3^Mut^ in a strain lacking endogenous Pex3 does not rescue the import of BFP-SKL into peroxisomes, confirming that this mutation disrupts the interaction (Figure 4J). Figures 4K - M contain representative microscopy images and the corresponding quantification showing that overexpression of this mutant in a background containing endogenous levels of wtPex3 to have functional peroxisomes, causes the same morphological phenotype as the overexpression of wtPex3. Thus, the interaction with Pex19 is not required for Pex3 to induce clustering of peroxisomes and lipid droplets.

### The triacylglycerol lipase Tgl4 is involved in the formation of the peroxisome-lipid droplet contact site

We sought to identify additional molecular players involved in the formation of contact sites upon Pex3 overexpression. Since many contact site tether proteins are involved in the formation of more than one contact site (Elbaz-Alon et al., 2015; Kvam & Goldfarb, 2004; Levine & Munro, 2001; Liu et al., 2017; Loewen et al., 2003; Murley et al., 2015; Toulmay & Prinz, 2012), we tested the involvement of known tether proteins of lipid droplets or peroxisomes with other organelles. Deletion of the peroxisomal-mitochondrial tethers Pex34 or Fzo1 (Shai et al 2018) did not affect formation of Pex3-dependent peroxisome and lipid droplet clusters (Supplemental Figure 3A), indicating that they are not required for formation of the contact sites. We also tested the involvement of the the splicing-generated pair of proteins Ldo16-Ldo45, which tether lipid droplets to the vacuole (Álvarez-Guerra et al., 2024; Diep et al., 2024). Deletion of these genes did not affect the clustering of peroxisomes or lipid droplets, nor the proximity of the clustered structure to the vacuole (Supplemental Figure 3B). Finally, it was recently shown that the human protein M1 Spastin tethers lipid droplets to peroxisomes (Chang et al., 2019). We deleted the yeast homolog Sap1, but observed no lteration of the Pex3-dependent phenotype (Supplemental Figure 3C).

We next performed a genome-wide microscopy-based screen. We crossed a strain carrying *TEF1pr*-*PEX3* and the peroxisomal marker Pex14-mKate2 with a genome-wide collection of deletion mutants of non-essential genes (Giaever et al., 2002) and hypomorphic DAmP allele mutants of essential genes (Breslow et al., 2008) using an automated mating and sporulation procedure (Cohen & Schuldiner, 2011; Hin et al., 2006). The resulting mutant collection contains *TEF1pr*-Pex3, Pex14-2xmKate2 and each gene deleted or depleted (Figure 5A). We analyzed this collection by automated microscopy after labeling cells with Bodipy and CMAC, to stain lipid droplets and vacuoles, respectively. The resulting images were analyzed manually searching for strains in which the phenotype was disrupted. We found a single hit in which the phenotype was significantly disrupted, which carried the deletions of the gene encoding for the lipase Tgl4 (Rajakumari et al., 2010).

**Figure 5:**
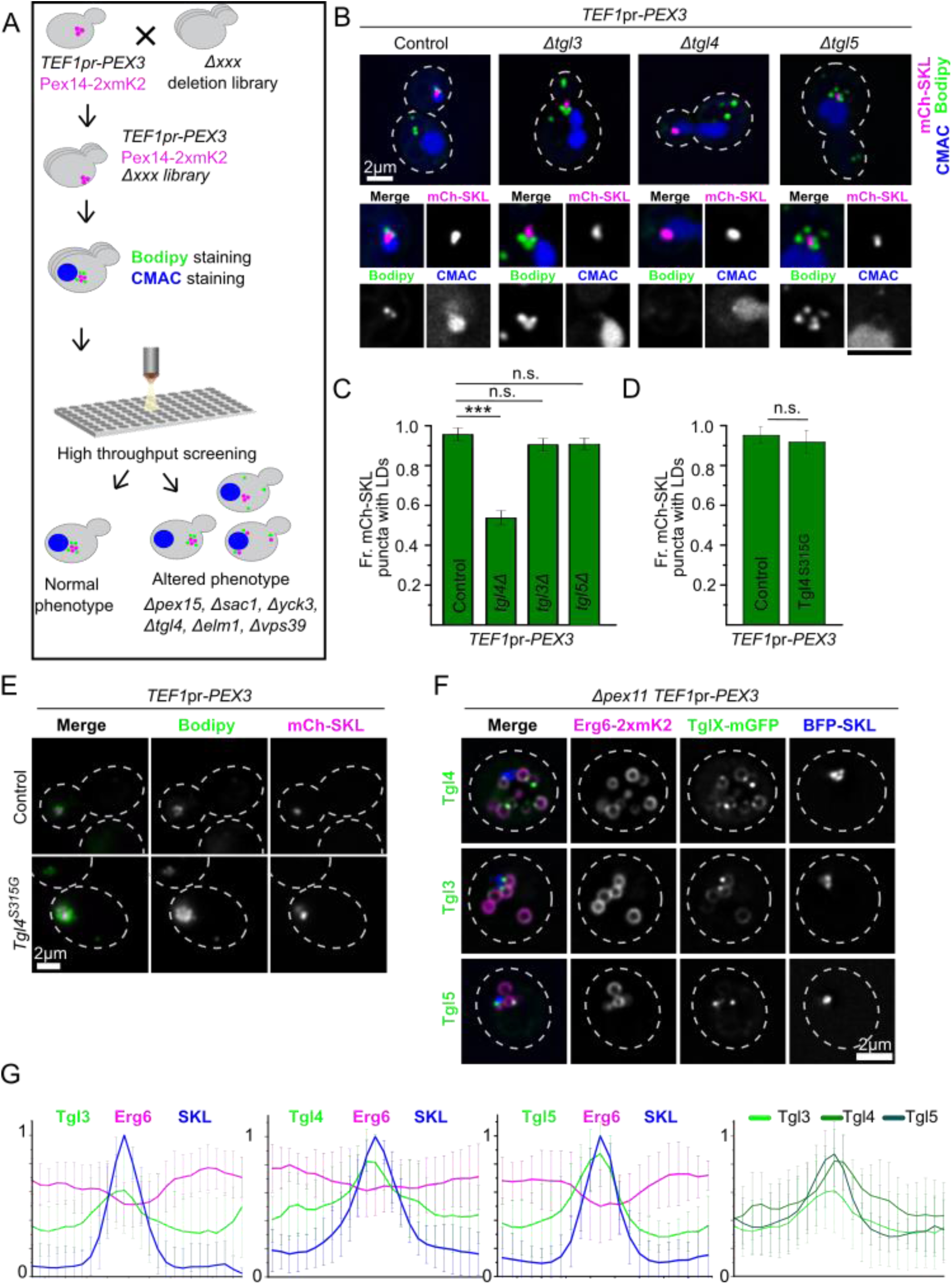
A genome-wide microscopy-based screen identifies that deletion of Tgl4 interferes with the targeting of LDs to the peroxisomal structure. A) A microscopy-based screen for deletions that disrupt the Pex3 overexpression phenotype. Establishment of a deletion library with the overexpression of Pex3 (TEF1pr-PEX3) by SGA and subsequent automated high-content microscopy-based screen. Deletion of TGL4 was identified to disrupt the phenotype. B-C) Deletion of TGL4 disrupts Per-LD CSs, and this effect is specific for this lipase. Panel B shows representative images of strains overexpressing Pex3 (TEF1pr-PEX3) in control cells and strains lacking either Tgl3, Tgl4 or Tgl5. All strains express mCherry-SKL construct to visualize the lumen of the peroxisomes, lipid droplets were stained with Bodipy and the vacuolar lumen was stained with CMAC. Cell outlines are shown as white dashed lines. Scale bars: 2 μm. Panel C shows the quantification of the proportion of peroxisomal structures with accumulations of lipid droplets next to them in the microscopy experiments described above. Three independent experiments were performed and 30 cells were analyzed for each experiment and condition. The different strains were compared by ANOVA and a post-hoc Tukey test. n.s., not significant, *** P < 0.001. D-E) The role of Tgl4 in establishing this contact site is independent from its lipase activity. Panel E shows representative images of strains overexpressing Pex3 (TEF1pr-PEX3) in control cells and in a strain with Tgl4 (S315G) punctual mutant in the background. Strains express BFP-SKL construct to visualize the lumen of the peroxisomes and Erg6 fused to 2xmKate2 to visualize the lipid droplets. Cell outlines are shown as white dashed lines. Scale bars: 2 μm. Panel D shows the quantification of the proportion of peroxisomal structures with accumulations of lipid droplets next to them. Three independent experiments were performed and 30 cells were analyzed for each experiment and condition. The two different strains were compared using an unpaired two-tailed Student’s t-test. n.s., not significant. F-G) Enlarged peroxisomes and LDs reveal that Tgl3, Tlg4 and Tgl5 are enriched at the interfaces at Per-LD CSs to different degrees. Representative pictures of a strain with overexpressed Pex3 (TEF1pr-PEX3), expressing the BFP-SKL construct to visualize the lumen of the peroxisomes, Erg6-2xmKate2 marking the lipid droplet monolayer, and each of the Tgl proteins tagged with mGFP. To produce enlarged peroxisomes and lipid droplets, the cells contain a deletion of PEX11, and were grown with oleate as the sole carbon source for 20hs. Cell outlines are shown as white dashed lines. Scale bars: 2 μm. The panels in G, show average line profiles of the BFP-SKL, Erg6-2xmKate and TglX-GFP signal around lipid droplets. The lines started at the opposite end of the peroxisome contact site, and aligned using the maxima of the BFP-SKL signal. 10 Cells were averaged for each strain. The final panel shows an overlay of the average line profiles of the different Tgl lipases, to compare the relative intensities.

This strain was manually re-constructed to confirm that the disruption of the phenotype did not depend on the genetic background used for the screen and re-analyzed by microscopy (Figure 5B). In cells that overexpressed Pex3, 95% of the peroxisomal structures were proximal to lipid droplets, while this number dropped to 51% in cells that in addition lacked Tgl4 (Figure 5C). Re-insertion of the *TGL4* ORF with its endogenous promoter in a plasmid fully recovered the interaction between peroxisomes and lipid droplets, indicating that the effect is specific to the lack of the gene, and not a secondary effect of the genomic modification (Supplemental Figure 4A and B).

We tested the specificity of the phenotype by deleting other lipid droplet-localized lipases. Neither deletion of the genes encoding for the lipases homologous to Tgl4, namely Tgl3 and Tgl5 (Athenstaedt & Daum, 2003, 2005) (Figure 5B and C) nor other ones, Tgl1, Ldh1 or Yeh1 (Jandrositz et al., 2005; Köffel et al., 2005; Thoms et al., 2011) (Supplemental Figure 4C and D) caused a disruption of the phenotype. The homologous lipases also did not cause a further disruption when combined with the deletion of *TGL4* (Supplemental Figure 4E and F). This suggests that it is the physical presence of the protein Tgl4 that affects the formation of the contact site and not its activity as a lipase. To test this hypothesis, we assessed the effect of mutation S315G, which disrupts its active site (Kurat et al., 2006), and observed no reduction in the association of lipid droplets to the peroxisomal cluster (Figure 5D and E).

In addition, we analyzed the localization of Tgl3, 4, and 5 on the lipid droplet surface upon induction of these contact sites. Again, we used the strain lacking *PEX11* and grew the cells in the presence of oleate to increase the size of peroxisomes and lipid droplets and gain spatial resolution. We observed that all three lipases were enriched in the region of the lipid droplet that was in contact with the peroxisomes in some cells, as can be appreciated in the example images and line profiles (Figure 5F and G). These enrichments, however, were not equally frequent for all lipases: Tgl4 and Tgl5 were enriched more frequently than Tgl3. This can be observed by the resulting peaks formed by averaging line profiles across many cells (Figure 5G).

To explore the requirements for the formation of Tgl4 foci at lipid droplet-peroxisome interfaces, we performed a microscopy-based screen (Supplemental Figure 5A). A *PEX3* overexpression allele, genes for lipid droplet and peroxisome visualization (Erg6-mCh and BFP-SKL), and Tgl4-GFP were introduced into the genome-wide deletion and DAmP libraries (Breslow et al., 2008; Giaever et al., 2002) by an automated mating approach (Cohen & Schuldiner, 2011; Tong and Boone 2006). Cells were cultured in the presence of oleate to expand lipid droplets and analyzed by automated microscopy. We identified a total of 86 mutants in which the accumulation of Tgl4-GFP foci at lipid droplet-peroxisome interfaces was fully or partially blocked (Supplemental Table 5, example microscopy images in Supplemental Figure 5B). We analyzed the common functions among the genes identified by the screen (Supplemental Figure 5C). The biggest group of genes was related to energy metabolism. This is also evidenced by the enrichment of the GO Terms “mitochondrion organization”, “mitochondrial respiratory chain complex assembly” and “mitochondrion” in our hit list with adjusted p-values of 0.0009, 0.001, and 0.008, respectively. Other groups of hits corresponded to hypoxia signaling, autophagy, and lipid homeostasis. Taken together, this suggests that the presence of Tgl4 at this organelle interface is regulated by the metabolic state of the cell. Additionally, we identified eight genes from the membrane contact site database, representing a 3-fold enrichment to the expected amount given the fraction of the genome annotated, likely reflecting the tight interrelations within the cellular contact site network.

### The phenotype of overexpression of Pex3 is conserved in metazoans

Since Pex3 is a conserved protein, and many of its functions, like the incorporation of peroxisomal membrane proteins and the role in pexophagy are conserved in metazoans, we decided to test if the phenotype of induction of contact sites is also conserved. Thus, we overexpressed Pex3-HA in *Drosophila melanogaster* larvae in the midgut using the GAL/UAS system (Brand & Perrimon, 1993). We then we observed cells from the midgut by fluorescence microscopy, comparing them to cells expressing only GFP-SKL, to observe peroxisomes when Pex3 is not overexpressed (Figure 6A and B). We observed that overexpression of Pex3 caused a reduction of the number of peroxisomes, as well as an increase in their size, whereas the number of lipid droplets was unaffected (Figure 6B - D). This phenotype was exacerbated during development, with a drastic increase in peroxisomal and lipid droplet size in the adult midgut (Figure 6E - H). We expressed Pex3-HA with another peroxisomal marker, YFP-SKL. Using Airyscan confocal microscopy, we found that Pex3-HA clearly labeled the surface of the enlarged peroxisome, while YFP was imported into the peroxisomal lumen by the peroxisomal targeting sequence, as expected (Figure 6I). Furthermore, these enlarged peroxisomes were closely associated with lipid droplets as can be observed by the shape deformation of the organelles when they are next to each other (Figure 6I). Thus, the phenotype closely resembles the one observed in yeast.

**Figure 6:**
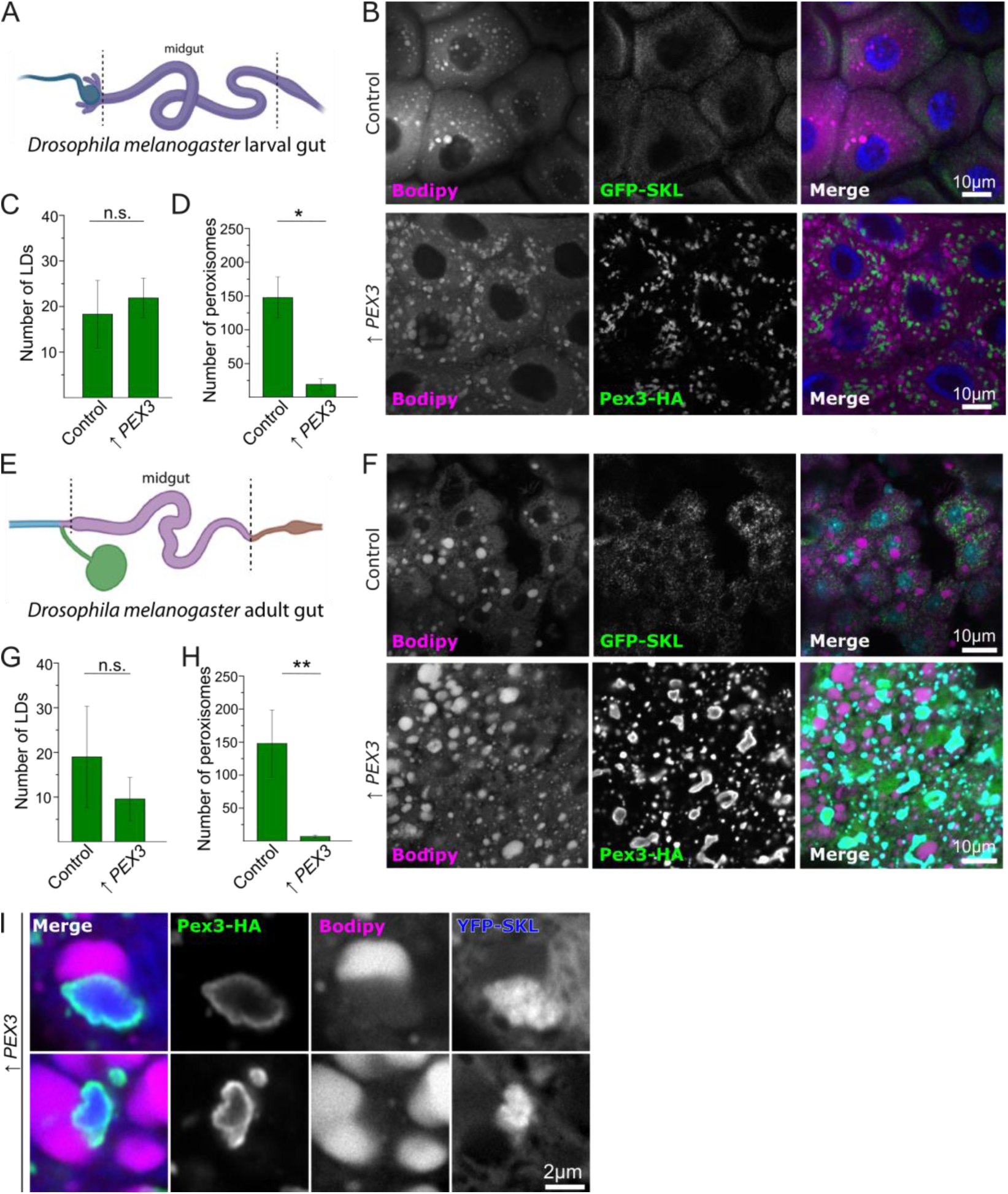
Overexpression of Pex3 leads to morphological changes in lipid droplets and peroxisomes in Drosophila melanogaster. A, E) Diagrams of the cuts from Drosophila melanogaster that were analyzed. Larval midgut (A) and midgut from adult flies (B) were analyzed. B-D) Representative images of control cells and strains overexpressing Pex3 in larval gut. Control cells express the GFP-SKL construct to visualize the lumen of the peroxisomes, while cells with overexpressed Pex3 also contain a GFP tag. Lipid droplets were stained with Bodipy and the nucleus with DAPI. Scale bars: 20 μm. Panels C and D show the quantification of the number of LDs (C) and peroxisomes (D). F-H) Representative images of cells from the adult midgut from control flies and flies overexpressing Pex3. Control cells express the GFP-SKL construct to visualize the lumen of the peroxisomes, while cells with overexpressed Pex3 also contain a GFP tag. Lipid droplets were stained with Bodipy and the nucleus with DAPI. Scale bars: 20 μm. Panels G and H show the quantification of the number of LDs (C) and peroxisomes (D). I) Representative Images of cells from adult flies midgut overexpressing Pex3-HA. Cells also express YFP-SKL to label the lumen of peroxisomes and lipid droplets were labeled with Bodipy. The enlarged organelles are observed in close proximity, with complementary morphological changes, suggesting the formation of membrane contact sites.

## DISCUSSION

Overexpression of Pex3 caused clustering of peroxisomes involving peroxisome-peroxisome contacts. The formation of these homotypic contact sites is an interesting phenomenon that poses the question of the functionality of contacts among the same type of organelles. For many organelles, the existence of specialized subpopulations has been proposed. The formation of homotypic contacts could be related to this, and represent contacts enabling communication among different specialized subopulations of one organelle. In this case, they would be functionally analogous to heterotypic contact sites, facilitating communication between distinct compartments. Importantly, homotypic peroxisome contacts have been observed before in cultured mammalian cells (Bonekamp et al., 2012; Schrader et al., 2000) and in tissues, even large-scale aggregates seem to be present (Gorgas & Zaar, 1984). Interestingly, homotypic contact sites have, to our knowledge, not been observed for most organelles. This could be related to the fact that, while most organelles can undergo homotypic fusion, it is thought that mature peroxisomes cannot fuse with each other (Mast et al., 2020). Thus, homotypic contact sites could serve as a mechanism to re-organize the separation of functions among specialized subtypes, a process that in other organelles could be achieved by fusion.

Another interesting aspect is the ever-growing multi-functional nature of the Pex3 protein. This is another example of membrane contact site tethers being multi-functional proteins. It is a recurring observation that organelles use some proteins as landmarks, which are recognized and bound by several different proteins which require interaction with that membrane. The most canonical example is the VAP-family protein in the endoplasmic reticulum (Loewen et al., 2003), but this phenomenon is also apparent for other organelles, like Vac8 for the vacuole and Tom70 for mitochondria (Álvarez-Guerra et al., 2024; Diep et al., 2024; Filadi et al., 2018; Hollenstein et al., 2019; Murley et al., 2015; Pan et al., 2000). These last two examples, alike Pex3, are involved in different processes in addition to contact site tethering (Schmidt et al., 2010; Veit et al., 2001; Wang et al., 1998). These multi-function proteins could aid the coordination or cross-regulation of different processes involving the same organelle. For example, the protein Vps39 is part of contact sites as well as a subunit of tethering complexes involved in vesicular transport, and the dynamics of the tethering complexes are linked to contact site formation (González Montoro et al., 2021).

This phenomenon could also explain the difference between the overexpression phenotype observed in *H. polymorpha* and in *S. cerevisiae*. Pex3 might have acquired its multiple functions sequentially in evolution, apparently resulting in the formation of different Pex3-dependent contact sites in different species. We also observed that the peroxisomal structure formed upon Pex3 overexpression was often (55% of the cases) in proximity to the vacuole, but the phenotype was partial, prompting us to study the more penetrant phenotypes in peroxisome and lipid droplet clustering. The fraction of the peroxisomal-lipid droplet clusters found next to the vacuole was not affected by deletion of the known lipid droplet-vacuole tethers Ldo16/45, indicating that this proximity is either generated by a parallel tethering complex, or depends on peroxisome tethering to the vacuole.

We also observed that Pex3 overexpression induced lipid droplet-peroxisomal contacts. These contacts have been observed in wild-type cells during growth in oleate (Binns et al., 2006), and occasionally included protrusions of the peroxisomes into the lipid droplets, that were positive for the β-oxidation enzyme Pox1. The proposed role for this contact site is an optimization of lipolysis-derived transfer of fatty acids for β-oxidation. Our finding of the lipase Tgl4 as a possible tether is very much in line with this. We observe that upon deletion of *TGL4*, the abundance of Pex3 overexpression-dependent peroxisome-lipid droplet contact sites decreases approximately by half. This suggests that tethering between these organelles does not exclusively depend on Tgl4, and the responsible tethers are yet to be described. In addition, Pex3 overexpression provides now a tool to massively induce these contact sites, which will aid their study. Recently, in mammalian cells the hereditary spastic paraplegia protein M1 Spastin was shown to form a tethering complex between lipid droplets and peroxisomes through its interaction with the transporter ABCD1. This contact is thought to aid in the trafficking of fatty acids through a mechanism that involves membrane deformation by ESCRT-III (Chang et al., 2019). The Tgl4-dependent contact site that we observe could have a different mechanism, and aid in fatty acid trafficking by local production of free fatty acids, as we observe several lipases to be enriched in the interface. In the future, it will be interesting to see how these different mechanisms interplay with each other, and to clarify if they operate in parallel in the same contact site, in structurally distinct ones, or in different species.

## MATERIALS AND METHODS

### Strains and plasmids

*Saccharomyces cerevisiae* strains were based on either BY4741 or W303. Genetic manipulations were carried out by homologous recombination of PCR-amplified cassettes as described in Janke et al. (2004). The Tgl4(S315G) mutant was generated by using CRISPR-Cas9 as described in Generoso (2016). For this, 500 ng of the plasmid pAGM218 and the oligonucleotide oAGM875 were transformed in the BY4741 strain. Plasmid pAGM218 was generated by PCR amplification of plasmid pCU5003 using primers oAGM873-874, and was confirmed by sequencing. We confirmed that the strain (AGMY1888) contained the appropriate mutation and no additional ones, by extracting genomic DNA, amplifying the region encoding for Tgl4 with primers oAGM803-804 and sequencing. All yeast strains used in this study are listed in Table S1.

The plasmid pAGM197 (pRS403 TEF1pr-BFP-SKL) was generated by Gibson Assembly, using primers oAGM839-840 and oAGM841-842 to amplify the BFP from plasmid CU5258 followed by an SKL tag, and the pRS403 backbone respectively. Purified PCR fragments were incubated with the Gibson Assembly master mix (New England Biolabs) for one hour at 55°C. The resulting plasmid was transformed in competent *E. Coli* DH5α and clones were checked by sequencing.

The cytosolic domain of Pex3 was amplified from genomic DNA from a BY4741 strain, using primers oAGM470-471. The purified DNA fragment was digested with NotI and SacII enzymes and ligated in a pRS416 vector containing a TEF1 promoter and a N-terminal GFP tag, which was digested the same way. After this plasmid was confirmed by sequencing, it was digested with XhoI and SacI to ligate the fragment corresponding to TEF1pr-GFP-Pex3(CD) into a pRS403 backbone, to generate plasmid pAGM134 (pRS403 TEF1-GFP-Pex3(CD)).

The plasmid pAGM212 (pRS403 TEF1pr-AlfaTag-mKate2-Pex3(CD)) was done by Gibson Assembly. Primers oAGM748-749 were designed to amplify the cassette TEF1pr-GFP-Pex3(CD) from plasmid pAGM134 and adding an Alfa tag at the N-terminus. The purified DNA was incubated with a fragment containing the mKate2 sequence (amplified from pAGM042 with primers oAGM750-751) and the Gibson Assembly master mix (New England Biolabs) for one hour at 55°C. The resulting plasmid was transformed in competent *E. Coli* DH5α and clones were checked by sequencing.

The plasmid pAGM210 (pRS403 TEF1pr-Pex3-mKate2) was also generated by Gibson Assembly. To obtain the corresponding fragments, primers oAGM851-852 were used to amplify the backbone (pRS403 TEF1pr) from plasmid pAGM134. Pex3 sequence was amplified from genomic DNA from a BY4741 strain with primers oAGM853-854. And last, mKate2 sequence was amplified by PCR using primers oAL855-856. All fragments were incubated with the Gibson Assembly master mix (New England Biolabs) for one hour at 55°C. The resulting plasmid was transformed in competent *E. Coli* DH5α and clones were confirmed by sequencing.

The plasmid pAGM211 (pRS403 TEF1pr-Pex3 (W128K, L131K)-mKate2) was generated by PCR amplification of the entire plasmid pAGM210, using primers oAL831-832 which contain the desired punctual mutations. The presence of these mutations was confirmed by sequencing.

C-terminal Alfa tag was added to pAGM210 by PCR amplification of the plasmid. For this, primers oAGM910-911 were used, which include the sequence of the ALFA tag, to generate plasmid pAGM225 (pRS403 TEF1pr-Pex3-mKate2-Alfatag). pAGM226 (pRS403 TEF1pr-Pex3 (W128K, L131K)-mKate2-Alfatag) was generated in the same way but using plasmid pAGM211 as a source instead. Both plasmids were confirmed by sequencing. All pRS403-based plasmids were integrated into the HIS3 locus.

Plasmid pRS315-mCherry-SKL was a gift from Judith Müller (Institute of Molecular Genetics and Cell Biology, Ulm University) and PRS415-GFP-SKL from (Schäfer et al., 2004). To obtain plasmid pAGM224, plasmid pRS315 GFP-SKL and pAGM197 were digested with SacI and XhoI. The purified fragment corresponding to TEF1pr-BFP-SKL was ligated into the pRS315 backbone. To stably integrate these constructs into the LEU2 locus, yeast transformations were done with DNA fragments amplified by PCR using primers oAGM090-091.

Plasmid pRS415 TGL4pr-Tgl4-msGFP2 was digested with ApaI and NotI restriction enzymes, taking the TGL4pr-Tgl4-msGFP2 construct. The purified DNA fragment was ligated to a pRS403 vector digested the same way. Plasmid pAGM242 (pRS403 TGL4pr-Tgl4-msGFP2) was confirmed by sequencing with oligonucleotide oAGM930.

All plasmids and oligonucleotides used in this study are listed in Table S1.

### Fluorescence microscopy, image quantifications and statistical analysis

Cells were grown to logarithmic phase in yeast extract peptone medium containing glucose (YPD), or synthetic medium supplemented with essential amino acids (SDC). For experiments with enlarged Lipid Droplets and peroxisomes, cells were grown in SC-Oleate media as described in Yifrach et al., 2022, and washed with SC media before imaging. The lumen of the vacuole was stained by adding 20 µM 7-amino-4-chloromethylcoumarin (CMAC) dye (Invitrogen) to 0.5 OD Units and incubating them for 15 min at 30°C with shaking, followed by one washing step in SDC medium. Proteins tagged with the HaloTag were labelled with the Janelia Fluor JFX650 ligand (Lavis Lab; (Grimm et al., 2021)). Yeast cells (0.5 OD Units) were incubated with 1.6 µM of JFX650 for 15 min, followed by eight washing steps with SDC medium (Day et al., 2018). Lipid droplets were stained with Bodipy dye (Echelon Biosciences, through MoBiTec). Cells were incubated with 1 µg/ml of the dye for 15 minutes at 30°C with shaking, followed by one wash with SDC medium.

In the majority of experiments, cells were imaged live in SDC medium on an Olympus IX-71 inverted microscope equipped with 100× NA 1.49 and 60× NA 1.40 objectives, an sCMOS camera (PCO, Kelheim, Germany), an InsightSSI illumination system, 4′,6-diamidino-2-phenylindole (DAPI), GFP, mCherry and Cy5 filters, and SoftWoRx software (Applied Precision, Issaquah, WA, USA). We used z-stacks with 200, 250 or 350 nm spacing for constrained-iterative deconvolution with the SoftWoRx software. For microscopy experiments of Tgl4 S315G mutant (Figure 5E), cells were imaged on a Olympus microscope IX-71 equipped with 100x oil NA=1.45, 60x oil NA=1.42 and 60x water NA= 1.2 objectives, a back-illuminated sCMOS camera (Hamamatsu ORCA Fusion-BT), a LED module (pe-800, CoolLED) and five different polychroic filter sets. The microscope is operated by Micro-Manager software version 2.0.3 nightly build (www.micro-manager.org). Images were deconvolved with Huygens Professional version 3.7.1 (Scientific Volume Imaging, The Netherlands, http://svi.nl).

All further image processing and quantification were performed using ImageJ (National Institutes of Health, Bethesda, MD, USA). One plane of the z-stack is shown in the figures.

The localization of proteins at interfaces of contact sites (Figures 2A, 2B and 5F) was quantified by performing a line profile around the lipid droplets observed in contact with a peroxisome. The intensity of signal in all channels was measured and values were manually aligned to the peak of the blue channel, corresponding to the BFP-SKL signal. Values were normalized to their maximum peak. For plots of Figures 2A and 2B, values of the representative images are shown. For plots in Figure 5G, averages and SD of ten cells (one peroxisome-lipid droplet interfase each) were plotted.

The colocalization measurement of the cytosolic domain of Pex3 with Lipid Droplets and peroxisomes (Figure 3C) was performed by applying a maximum entropy threshold to all channels using Image J. Cells were manually defined as regions of interest (ROIs) and the Manders M1 and M2 coefficients for each cells were obtained with the JACoP plugin for ImageJ (S. Bolte & F. P. Cordelieres, 2006)

For the distances measurement between the peroxisomes and mitochondria in figure 3F, the far red and green channels were analyzed. An Otsu threshold was applied to the raw data and 3D ROIs corresponding to peroxisomes and mitochondria were built with the 3D ROI manager plugin from the 3D ImageJ Suite and manually curated (Ollion et al., 2013). Border to border distances were measured using the 3D ROI manager between the different objects (peroxisomes and mitochondria) and the shortest distances for each peroxisome to mitochondria was plotted.

### TurboID assay and mass spectrometry

To assess the molecular environment of overexpressed Pex3 by TurboID, three cultures of each strain, AGMY192 and AGMY575, were grown up to logarithmic phase in YPAD media. 100 µM of Biotin (Novabiochem) was added to the cultures and incubated for 3 hours. 350 OD units were harvested, washed with sterile water and split into two vials. Cells were resuspended in 650 µl lysis buffer (20 mM TrisHCl pH 8, 1% sodium dedecilsulfate (SDS), 1 mM dithiotreitol (DTT)) and lysed twice in a Fastprep machine (MP Biomedicals) at 40 m/s for 40 seconds. Samples were treated with a heat shock (60°C for 10 minutes) and after cooling down, 650 µl of buffer B were added (20 mM TrisHCl pH 8, 0,5% SDS, 8 M Urea, 1 mM DTT). Lysates were centrifuged at 20.000 g for 20 minutes. Supernatants belonging to the same sample were combined and protein concentration was measured using Bradford assay (Bio-Rad). 100 µl of slurry Streptavidin agarose beads (Thermo Fisher Scientific) were equilibrated for each sample which were added in equal protein concentrations. The samples were incubated for one hour at room temperature with rotation. The beads were centrifuged at 300 × g for 1 min and washed four times with 1 ml wash buffer (20 mM TrisHCl pH 8, 0,75% SDS, 4 M Urea, 1 mM DTT) and four times with the same buffer without detergent. The samples were processed for mass spectrometry with the iST 96x Kit (Preomics) according to manufacturer instructions.

For mass spectrometry analysis, reversed-phase chromatography was performed on a Thermo Ultimate 3000 RSLCnano system connected to a Q-ExactivePlus mass spectrometer (Thermo Fisher Scientific) through a nano-electrospray ion source. For peptide separation, 50 cm PepMap C18 easy spray columns (Thermo Fisher Scientific) with an inner diameter of 75 µm were used and kept at a temperature of 40 °C. The peptides were eluted from the column with a linear gradient of acetonitrile from 10 to 35% in 0.1% formic acid for 118 min at a constant flow rate of 300 nl/min, followed by direct electrospraying into the mass spectrometer. The mass spectra were acquired on the Q-Exactive Plus in a data-dependent mode to automatically switch between full scan MS and up to ten data-dependent MS/MS scans. The maximum injection time for full scans was 50 ms, with a target value of 3,000,000 at a resolution of 70,000 at m/z = 200. The ten most intense multiply charged ions (z = 2) from the survey scan were selected with an isolation width of 1.6 Th and fragments with higher energy collision dissociation with normalized collision energies of 27 (Olsen et al., 2007). Target values for MS/MS were set at 100,000 with a maximum injection time of 80 ms at a resolution of 17,500 at m/z = 200. To avoid repetitive sequencing, the dynamic exclusion of sequenced peptides was set at 30 s.

The resulting MS and MS/MS spectra were analyzed using MaxQuant (version 1.6.0.13, https://www.maxquant.org/) utilizing the integrated ANDROMEDA search algorithms (Cox et al., 2011; Cox & Mann, 2008). The peak lists were compared against local databases for *S. cerevisiae* (obtained from the *Saccharomyces* Genome database, Stanford University), with common contaminants added. The search included carbamidomethylation of cysteine as a fixed modification and methionine oxidation, N-terminal acetylation, and phosphorylation as variable modifications. The maximum allowed mass deviation was 6 ppm for MS peaks and 20 ppm for MS/MS peaks. The maximum number of missed cleavages was two. The false discovery rate was 0.01 on both the peptide and the protein level. The minimum required peptide length was six residues. Proteins with at least two peptides (one of them unique) were considered identified. The re-quant option of MaxQuant was disabled. The calculations and plots were performed using the Perseus software (Tyanova et al., 2016)

### On-section CLEM tomography

#### High pressure freezing (HPF)

For high-pressure freezing, yeast cells were grown in YPD (1% yeast extract, 2% peptone, 2% glucose) medium to OD600 of 0.4-0.6. The suspension was concentrated by vacuum filtration onto a 0.45 µm membrane filter (Merck, HVWG04700), which was was placed onto an agar plate and the yeast paste was scraped using a pipette tip. Then, 3 µl of the concentrated paste were transferred into a 100 µm deep cavity of an aluminum planchette (Engineering Office M. Wohlwend GmbH, 241) until the cavity was overfilled. Subsequently, the flat side of a planchette (Engineering Office M. Wohlwend GmbH, 242) was placed on top and excess yeast paste was quickly removed. The finished assembly was immediately subjected to high-pressure freezing using a HPF Compact 03 (Engineering Office M. Wohlwend GmbH). Vitrified samples were stored in liquid nitrogen until further processing via freeze substitution.

#### Freeze substitution (FS)

Freeze substitution and Lowicryl embedding was performed as described in (Ader & Kukulski, 2017) with slight modifications. Briefly, samples were freeze substituted in 0.1% uranyl acetate (Science Services, E22400) in anhydrous acetone (VWR, 83683.230) for 24 h at -90 °C. Then, the temperature was raised to -45 °C (5°C/h) and the samples were washed three times with anhydrous acetone. Next, infiltration with increasing concentrations (10%, 25%, 50%, 75%) of Lowicryl HM20 in acetone (Science Services, PS14340) was carried out for 2 h each step. During the last two steps the temperature was raised by 10 °C each to -35 °C and -25 °C respectively. Afterwards 100% Lowicryl was exchanged three times in 10 h steps. Finally, polymerization was carried out via UV light for 24 h at -25 °C and further 24 h at 20 °C. Polymerized sample blocks were taken out of the AFS and were ready for ultrathin sectioning.

#### On-section light microscopy

For light microscopy imaging, 200 mesh carbon film grids (Plano, S160) containing 250 nm thin sections of the samples were placed on a 20 µl drop of PBS, pH 8, on a 25 mm coverslip. The grids were sandwiched between another 25 mm coverslip and transferred to a custom-made holder. Z-stacks were acquired using an Olympus FV-3000 operated as a wide-field setup, equipped with an sCMOS camera (ORCA-Flash 4.0, Hamamatsu, Japan) and a 60x oil immersion objective (PLAPON-SC NA 1.4). Overview images were taken with a 10x objective (UPL SAPO NA 0.4) to facilitate later correlation within the transmission electron microscopy (TEM).

#### TEM tomography acquisition, correlation and segmentation

For TEM tomography, sections were labelled on both sides with 10 nm protein A gold fiducials. Sections were then contrasted with 3% uranyl acetate for 30 min and 2% lead citrate for 20 min in the LEICA EM AC20 and subsequently analysed with a TEM at 200kV (JEM2100Plus, JEOL, Japan) equipped with a 20-megapixel EMSIS Xarosa CMOS camera (EMSIS, Germany). Regions of interest from the LM were relocated within the TEM and tilt series from +-65° with 1° increments were acquired using TEMography software (TEMography.com, JEOL, Japan). The tomograms were then reconstructed using the back projection algorithm in IMOD (Kremer, Mastronarde et al. 1996). The tomography data were manually overlaid with the fluorescence signal from the LM using Adobe Photoshop.

### Pulldowns and Western blot

Cells were grown to logarithmic phase in yeast extract peptone medium containing glucose (YPD) and 100 OD600 units were harveste by centrifugation. The pellets were resuspended in lysis buffer (50 mM HEPES pH 7,4, 150 mM NaCl, 10% glycerol, 1% CHAPS, 1 mM PMSF, protease inhibitor cocktail FY (Serva) 1/100) and lysed twice in a FastPrep device (6 m/s for 40 s; MP Biomedicals), with a 5-min incubation on ice in between. The lysate was centrifuged at 20.000 xg for 20 min and 4 °C. 12.5 µl slurry GFP-Trap agarose beads (Chromotek) or Alfa agarose beads (ALFA selector ST, NanoTag Biotechnologies) were equilibrated with lysis buffer and the same protein concentration was added to the beads. Samples were incubated at 4 °C for 15 min with rotation. The beads were centrifuged at 300 ×g for 1 min and washed twice with 1 ml lysis buffer and four times with 1 ml buffer without detergent. Proteins were eluted from the beads by incubating them with 1X Laemmli buffer (4% SDS, 0.05% bromophenol blue, 0.0625 M Tris, pH 7.4, 2.5% β-mercaptoethanol, and 10% glycerol) at 95°C for 10 minutes.

Proteins were separated using SDS-PAGE in 10% Bis-Tris acrylamide/ bisacrylamide gels and transferred to a nitrocellulose membrane (GE Healthcare). The membranes were blocked for 30 min with PBS 5% milk and incubated with the first antibody at 4 °C overnight with gentle shaking. The membranes were washed three times with PBS, and once with TBS-Tween (0.5% (v/v) Tween 20) for 5 minutes. A fluorescent-dye-coupled secondary antibody (Thermo Fisher Scientific) was diluted 1:20000 in PBS 5% milk and incubated at room temperature for 1 hour. A Bio-Rad ChemiDoc MP imaging system was used to detect the fluorescent signal. The whole Western is shown in Supplemental Figure 2A. Supplemental Figure 2B shows the right side of the membrane (previously decorated with antibodies against the Alfa tag) after stripping and decorated with antibodies against GFP. For stripping, the membrane was washed with distilled water, incubated for 5 minutes with 1M sodium hydroxide and washed again. Once the lack of fluorescent signal was confirmed by scanning, the membrane was blocked and incubated with the anti-GFP antibody as specified previously. The antibodies used are listed in Table S4.

### Fly husbandry

Flies were reared on standard cornmeal food (130g yarn agar, 248g Baker’s yeast, 1223g Cornmeal and 1.5 l sugar beet syrup in 20 l distilled water) and kept in a 25°C incubator with light-dark-cycle. Fly lines used in this study were mex-Gal4 (kindly provided by the lab of Irene Miguel-Aliaga), UAS-Pex3-GFP, UAS-Pex3-HA (kindly provided by the lab of Reinhard Bauer) and UAS-GFP-SKL (Bloomington Drosophila stock center #28882). For studies in the larval gut, the following genotypes were used: w; mex-Gal4; UAS-GFP-SKL and w; mex-Gal4; UAS-GFP-SKL/UAS-Pex3-HA. For studies in the adult gut, the following genotypes were used: w; mex-Gal4; UAS-GFP-SKL, w; mex-Gal4; UAS-GFP-SKL/UAS-Pex3-HA and w; mex-Gal4; UAS-YFP-PTS1/UAS-Pex3-HA. Animals were reared on standard diet and analyzed as 3rd instar larvae or 5 day old adults (male and female), respectively.

### Imaging of Drosophila guts

Antibodies used in this study were α-GFP (Santa Cruz Biotechnology) and α-HA (Invitrogen), α-TOMM20 (Sigma-Aldrich). For immunohistochemistry, guts from 3^rd^ instar larvae or adult flies were dissected in PBS and fixed for 1 h in 0.5 % PBS-Tween20 and 4 % formaldehyde. Tissue was washed with 0.5 % PBS-Tween20 and blocked with donkey serum before incubation with the primary antibody (overnight at 4 °C). The tissue was washed in 0.1 % PBS-Tween20 before incubation with BODIPY 581/591 (Thermo Scientific) and the secondary antibody at room temperature in the dark for 1 h. Secondary antibodies coupled to Alexa or Cyanine dyes were from Molecular probes. The tissue was washed and incubated for 5 min with DAPI (4’,6-diamidino-2-phenylindole). For imaging, we used a Zeiss LSM 710 with a 25x water lens (Plan-Neofluar, Zeiss), 40x water lens (C-Apochromat, Zeiss), and 63 x water lens (Plan-Apochromat, Zeiss) and a Zeiss LSM 880 with Airyscan detector. We used ImageJ to quantify peroxisome and lipid droplet number and area from at least 3 individual cells from different experiments.

### Library generation and high-throughput microscopy

All yeast manipulations were performed in high-density format (384–1,536 strains per plate) using a RoToR bench-top colony array instrument (Singer Instruments). In order to find key proteins involved in the formation of the peroxisome-peroxisome or peroxisome-lipid droplet contact sites (Figure 5A), strain AGMY1303, with an overexpression of Pex3 (*TEF1*pr-Pex3) and a peroxisomal marker (Pex14-2xmKate2) was crossed with a genome-wide library of deletion (Giaever et al., 2002) and hypomorphic allele (Breslow et al., 2008) strains, by the synthetic genetic array method (Cohen & Schuldiner, 2011; Hin et al., 2006). For analysis of Tgl4 localization (Supplemental Figure 5), strain yMB1326 (*TEF2*pr-Pex3 Erg6-mCh BFP-SKL Tgl4-GFP) was crossed with the same mutant collections.

Cells were mated on rich medium plates, diploids were selected and sporulation was induced by incubating the cells for five to eight days on nitrogen starvation media plates. Haploid cells were selected on 50 mg/L Canavanine and 50 mg/L Thialysine. Finally, haploid cells containing the combination of all desired manipulations were selected. A subset of strains was verified by microscopy and confirmed by PCR.

For the automated imaging of the obtained libraries, cells were first transferred from agar plates into 384 well plates for growth in liquid medium. For the screen for genes involved in formation of lipid droplet-peroxisome contacts (Figure 5A), cultures were grown overnight at 30°C in SDC. A JANUS liquid handler (PerkinElmer) connected to the incubator was used to dilute the strains to an OD600 of ∼0.2, and plates were incubated at 30°C for 4 h. Cells were washed and fresh SDC media was added containing 20 µM 7-amino-4-chloromethylcoumarin (CMAC) dye and 1 µg/ml of Bodipy dye, followed by half an hour incubation and another wash step. For the Tgl4-GFP screen (Supplemental Figure 5), liquid handling was performed using a MicroPro 300 liquid handler. Following the same overnight culture as the previous screen, cells were grown in the presence of 0.2% oleate in synthetic medium for 24 hours for lipid droplet enlargement.

Strains were then transferred by a liquid handler into glass-bottom 384-well microscope plates (Matrical Bioscience) coated with concanavalin A (Sigma-Aldrich) and incubated for 20 minutes to allow adhesion of cells to the bottom of the plates. Afterwards, wells were washed twice with SDC medium (SC medium for oleate treated cells) to remove non-adherent cells leaving a cell monolayer. Plates were then transferred to an Olympus automated inverted fluorescence microscope system. In the screen for lipid droplet-peroxisome contact site proteins (Figure 5A), cells were imaged in SDC at 18–20°C using a 60× air lens (NA 0.9) and with an ORCA-ER charge-coupled device camera (Hamamatsu), using ScanR software. In the Tgl4-GFP screen (Supplemental Figure 5), cells were imaged in SC medium at 18-20 °C using a 60× air lens (NA 0.9) and with an ORCA-flash4.0 camera (Hamamatsu), using ScanR software. After acquisition, images were manually reviewed using ImageJ (National Institutes of Health).

## AKNOWLEDGEMENTS

We thank Christian Wingen and Fatmire Bujupi for generating the UAS-Pex3-GFP and UAS-Pex3-HA fly lines. We thank Hadar Meyer and Yeynit Asraf for their assistance with the high-throughput screens.

This project was funded through a Deutsche Forschungsgemeinschaft (DFG) individual research grant to Ayelén González Montoro (GO3313/1-1). Work performed in the Bohnert lab was supported by the DFG, projects SFB1557 P3 and SFB1348 A13. Work in the Bülow lab was supported by DFG grants 417982926 and 535112684. Work on peroxisomes in the Schuldiner lab is supported by an Israel Science Foundation grant ISF 914/22. The robotic system of the Schuldiner lab was purchased through the kind support of the Blythe Brenden-Mann Foundation. MS is an Incumbent of the Dr. Gilbert Omenn and Martha Darling Professorial Chair in Molecular Genetics

## SUPPLEMENTAL FIGURES

**Supplemental Figure 1:**
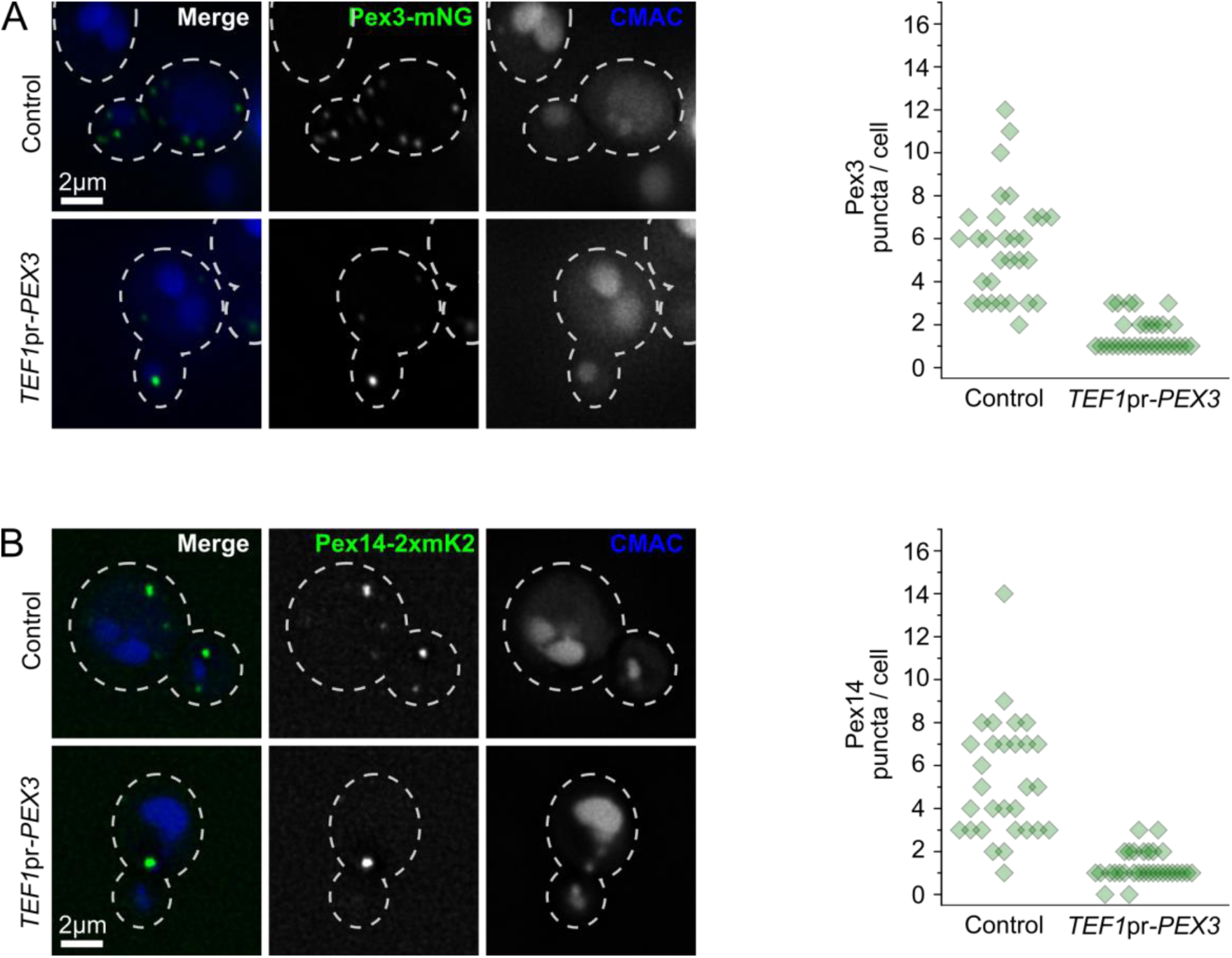
Overexpression of Pex3 reduces the number of puncta of different peroxisomal markers. **A)** Representative pictures of a strain expressing Pex3 fused to mNeonGreen to visualize the peroxisomes, either at endogenous levels (Control) or overexpressed (*TEF1*pr-*PEX3*) and the vacuolar lumen stained with CMAC. Cell outlines are shown as white dashed lines. Scale bar: 2 μm. The quantification of the amount of Pex3 puncta per cell is shown to the right. 30 cells from a single experiment were analyzed for each condition. Diamonds correspond to individual cells. **B)** Representative pictures of a strain expressing Pex14 fused to 2xmKate2 to visualize the peroxisomes, either with Pex3 at endogenous levels (Control) or overexpressed (*TEF1*pr-*PEX3*) and the vacuolar lumen stained with CMAC. Cell outlines are shown as white dashed lines. Scale bar: 2 μm. The quantification of the amount of Pex14 puncta per cell is shown to the right. 30 cells from a single experiment were analyzed for each condition. Diamonds correspond to individual cells.

**Supplemental Figure 2:**
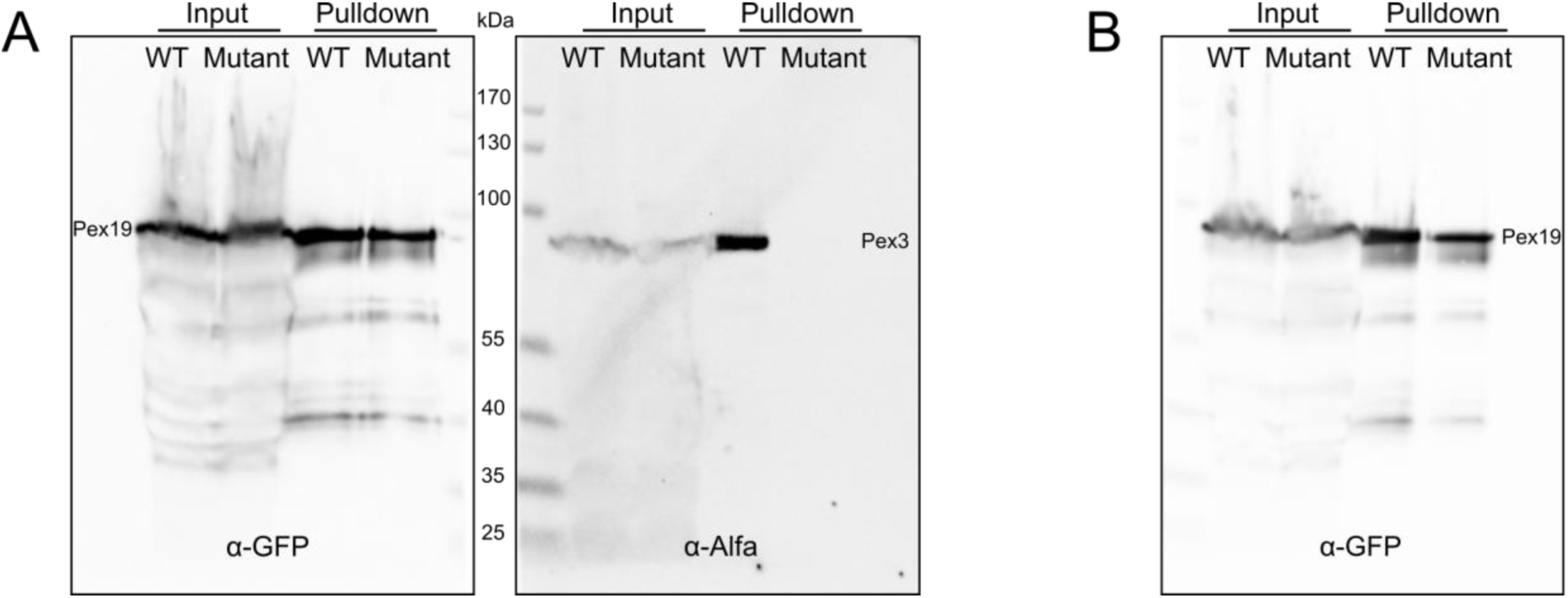
W128K and L131K mutations in Pex3 abolish interaction with Pex19. **A)** Whole Western blot from affinity purification of msGFP2-Pex19. The left side of the membrane was revealed against GFP and the right side against AlfaTag. Pex3-mKate2-AlfaTag co-purifies with msGFP2-Pex19 while Pex3(W128K L131K)-mKate2-AlfaTag does not. **B)** The right side of Western blot from panel A, previously revealed against AlfaTag, was stripped and revealed against GFP, showing comparable amounts of msGFP2-Pex19 purified for both strains.

**Supplemental Figure 3:**
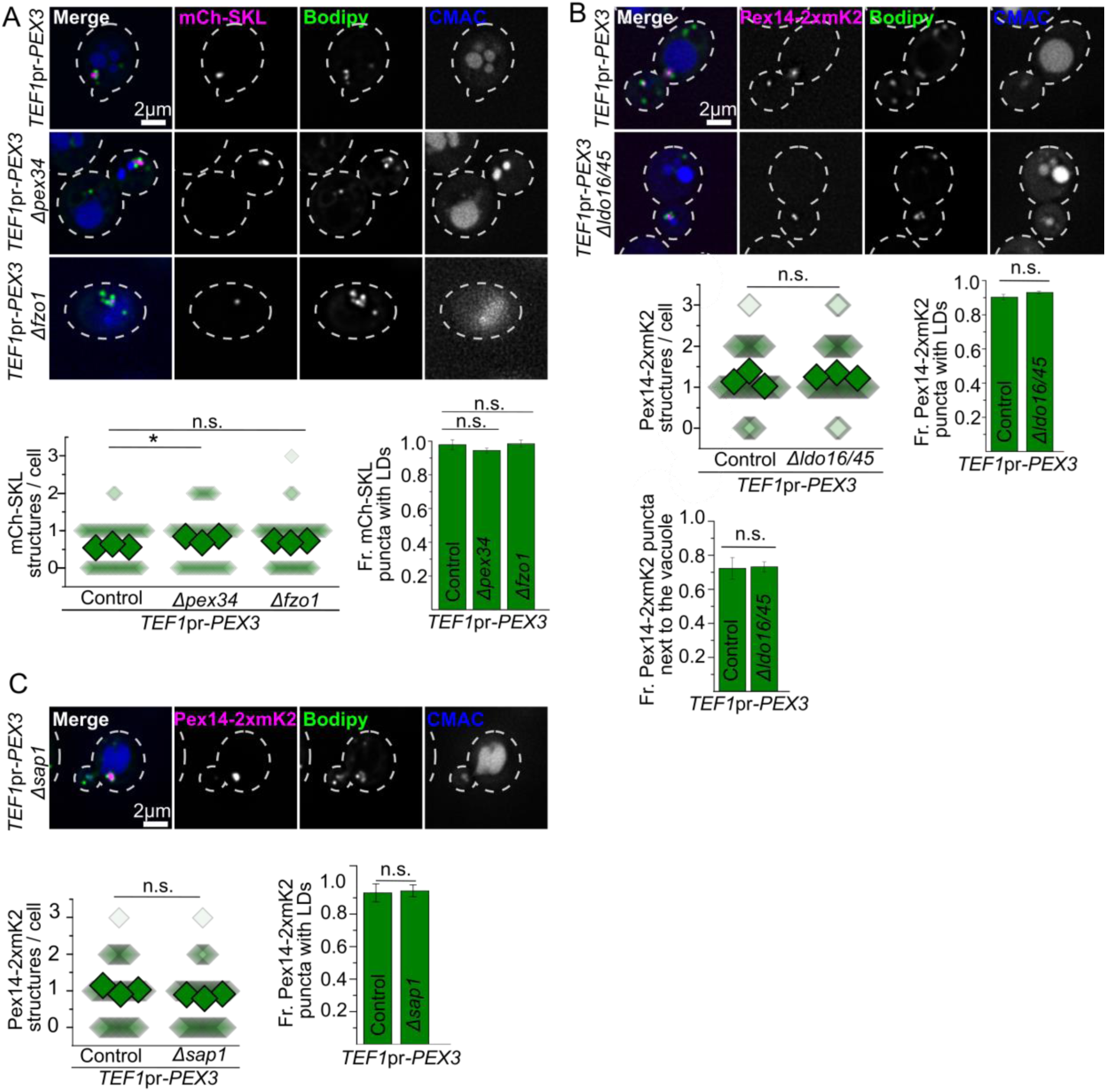
Formation of the structure is independent of known peroxisome or lipid droplet contact site tethers. A) Representative images of strains overexpressing Pex3 (TEF1pr-PEX3) in control cells and strains lacking Pex35 or Sap1. Strains express the mCherry-SKL construct to visualize the lumen of the peroxisomes, lipid droplets were stained with Bodipy and the vacuolar lumen was stained with CMAC. Cell outlines are shown as white dashed lines. Scale bars: 2 μm. Quantifications are shown below. For the quantification of the amount of peroxisomal structures per cell, small diamonds correspond to individual cells, bigger diamonds correspond to the average of independent experiments. Three independent experiments were performed and 30 cells were analyzed for each experiment and condition. Statistical differences were compared with ANOVA and a post-hoc Tukey test. n.s., not significant; * P < 0.05. B) Representative images of strains overexpressing Pex3 (TEF1pr-PEX3) in control cells and strains lacking Ldo16 and Ldo45. Pex14 is tagged with 2xmKate2 to visualize construct to visualize the peroxisomes, lipid droplets were stained with Bodipy and the vacuolar lumen was stained with CMAC. Cell outlines are shown as white dashed lines. Scale bars: 2 μm. Quantifications are shown below. For the quantification of the amount of peroxisomal structures per cell, small diamonds correspond to individual cells, bigger diamonds correspond to the average of independent experiments. Three independent experiments were performed and 30 cells were analyzed for each experiment and condition. Statistical differences were compared using an unpaired two-tailed Student’s t-test. n.s., not significant. C) Representative images of a strain overexpressing Pex3 (TEF1pr-PEX3) and lacking Sap1. Pex14 is tagged to visualize construct to visualize the peroxisomes, lipid droplets were stained with Bodipy and the vacuolar lumen was stained with CMAC. Cell outlines are shown as white dashed lines. Scale bars: 2 μm. Quantifications are shown below. For the quantification of the amount of peroxisomal structures per cell, small diamonds correspond to individual cells, bigger diamonds correspond to the average of independent experiments. Three independent experiments were performed and 30 cells were analyzed for each experiment and condition. Statistical differences were compared using an unpaired two-tailed Student’s t-test. n.s., not significant.

**Supplemental Figure 4:**
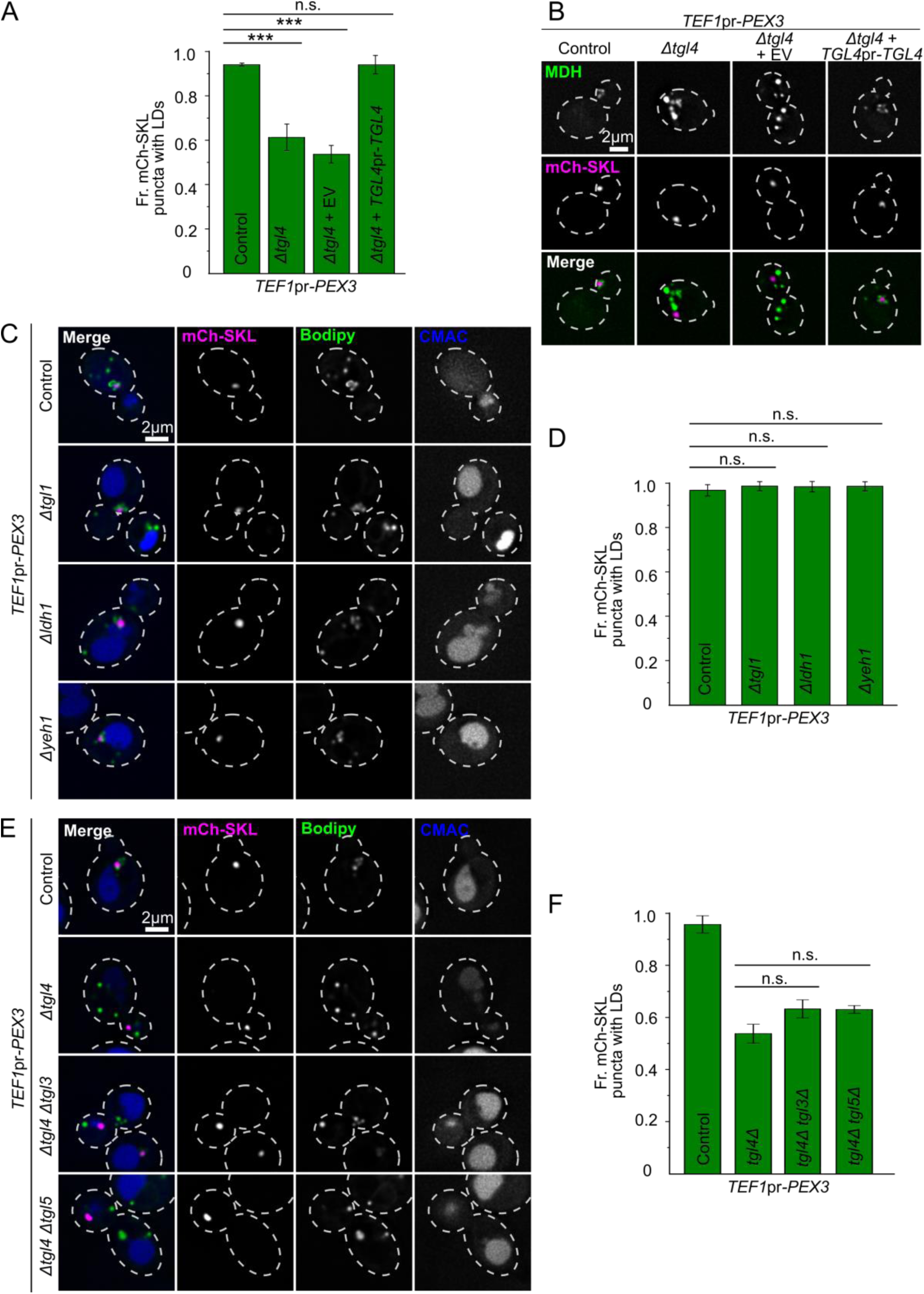
Disruption of the lipid droplet association to peroxisomes in Pex3 overexpression cells is specific for Tgl4 among lipases. **A – B)** Disruption of peroxisomes-lipid droplets interactions caused by *tgl4Δ* is rescued by re-introducing Tgl4 in a vector. Microscopy experiments were done comparing strains overexpressing Pex3 (TEF1pr-PEX3) in control cells, *tgl4Δ* and in *tgl4Δ* were either an empty vector or a vector containing Tgl4 ORF were introduced. Panel A shows the quantification of the fraction of peroxisomal structures with accumulations of lipid droplets next to them. Three independent experiments were performed and 30 cells were analyzed for each experiment and condition. The different strains were compared by ANOVA and a post-hoc Tukey test. n.s., not significant, *** P < 0.001. Panel B shows representative images of the microscopy experiment described above. All strains express mCherry-SKL construct to visualize the lumen of the peroxisomes, lipid droplets were stained with Bodipy and the vacuolar lumen was stained with CMAC. Cell outlines are shown as white dashed lines. Scale bars: 2 μm. **C – D)** Deletion of the lipases Tgl1, Ldh1 or Yeh1 has no effect on the phenotype of Pex3 overexpression. Panel C shows representative images of strains overexpressing Pex3 (*TEF1*pr-*PEX3*) in control cells, or *tgl1Δ*, *ldh1* or *yeh1Δ* cells. All strains express mCherry-SKL construct to visualize the lumen of the peroxisomes, lipid droplets were stained with Bodipy and the vacuolar lumen was stained with CMAC. Cell outlines are shown as white dashed lines. Scale bars: 2 μm. Panel D shows the quantification of the fraction of peroxisomal structures with accumulations of lipid droplets next to them in the microscopy experiments described above. Three independent experiments were performed and 30 cells were analyzed for each experiment and condition. The different strains were compared by ANOVA and a post-hoc Tukey test. n.s., not significant. **E – F)** Deletion of Tgl5 or Tgl3 in addition to Tgl4 has no additive effect on the Pex3 overexpression phenotype. Panel E shows representative images of strains overexpressing Pex3 (*TEF1*pr-*PEX3*) in control cells, or *tgl4Δ*, *tgl4Δtgl3Δ* or *tgl4Δtgl5Δ* cells. All strains express mCherry-SKL construct to visualize the lumen of the peroxisomes, lipid droplets were stained with Bodipy and the vacuolar lumen was stained with CMAC. Cell outlines are shown as white dashed lines. Scale bars: 2 μm. Panel F shows the quantification of the fraction of peroxisomal structures with accumulations of lipid droplets next to them in the microscopy experiments described above. Three independent experiments were performed and 30 cells were analyzed for each experiment and condition. The different strains were compared by ANOVA and a post-hoc Tukey test. n.s., not significant.

**Supplemental Figure 5:**
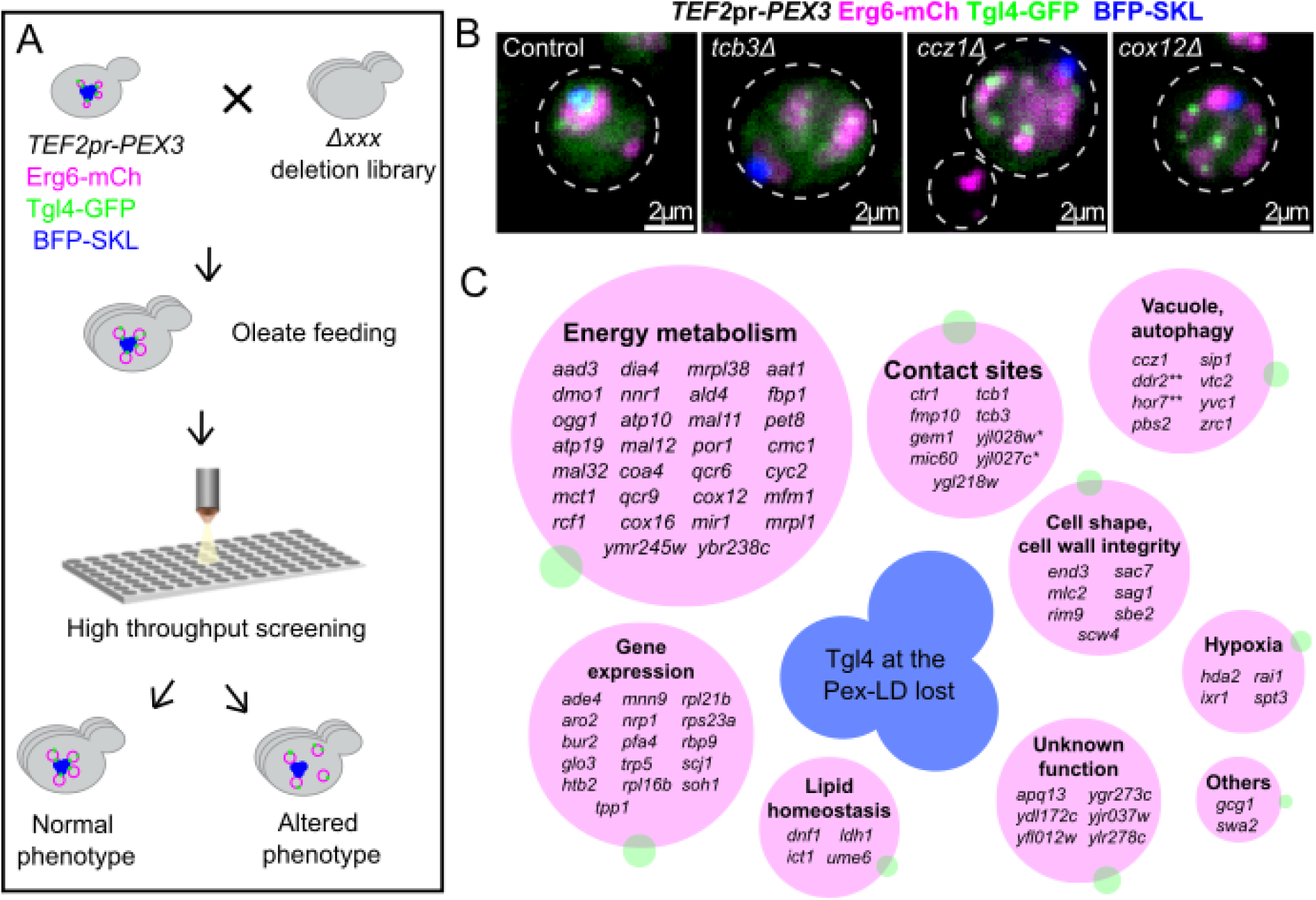
A genome-wide screen for mutations affecting the localization of Tgl4 to the peroxisome-lipid droplet interface. **A)** A microscopy-based screen for deletions/DAmP alleles that disrupt the enrichment of Tgl4 in the interface between peroxisomes and lipid droplets. Establishment of a deletion library with the overexpression of Pex3 (*TEF2pr-PEX3*), Erg6-mCherry, BFP-SKL and Tgl4-GFP by automated mating and subsequent automated high-content microscopy. **B)** Example microscopy images of three deletions (*tcb3Δ, ccz1Δ and cox12Δ*) among the 86 deletions that fully or partially disrupt the enrichment of Tgl4 to the peroxisome-lipid droplet interface. Scale bars: 2 μm. **C)** Aggrupation of the hits of the screen into functional categories based on their annotations in the *Saccharomyces* genome database. *, adjacent genes; **, paralogs.

**Supplemental Table 1:** *Saccharomyces cerevisiae* strains used in this study.

| **Name** | **Identifier** | **Genotype** | **Source** |
| --- | --- | --- | --- |
| BY4741 | AGMY002 | *MATa his3Δ1 leu2Δ0 met15Δ0 ura3Δ0* | Euroscarf |
| W303 | AGMY064 | *matAlpha ura3-52 trp1Δ 2 leu2-3112 his3-11 ade2-1 can1-100* | Ungermann Lab |
| TEF1pr-Pex3 | AGMY192 | *BY4741 – PEX3pr::TEF1pr-NatNT2* | This study |
| mCherry-SKL | AGMY1267 | *BY4741 – LEU2::mCherry-SKL-LEU2* | This study |
| TEF1pr-Pex3 mCherry-SKL | AGMY1268 | *BY4741 – PEX3pr::TEF1pr-NatNT2 LEU2::mCherry-SKL-LEU2* | This study |
| TEF1pr-Pex3-TurboID-V5 | AGMY575 | *BY4741 – PEX3pr::TEF1pr-NatNT2 PEX3::TurboID-V5-KanMX* | This study |
| ΔPex3 | AGMY1847 | *BY4741 – Pex3Δ::HphNT1* | This study |
| ΔPex11 | AGMY894 | *BY4741 – Pex11Δ::KanMX* | This study |
| ΔPex11 TEF1pr-Pex3-msGFP Erg6-2xmKate2 pRS403 TEF1pr-BFP-SKL | AGMY1809 | *BY4741 – Pex11Δ::KanMX PEX3pr::TEF1pr-NatNT2 ERG6::2xmKate2-URA3 HIS3::pRS403-TEF1pr-BFP-SKL-HIS3 PEX3::msGFP-HphNT1* | This study |
| ΔPex11 TEF1pr-Pex3 Erg6-2xmKate2 pRS403 TEFpr-BFP-SKL Pex13-msGFP | AGMY1824 | *BY4741 – Pex11Δ::KanMX PEX3pr::TEF1pr-NatNT2 ERG6::2xmKate2-URA3 HIS3::pRS403-TEF1pr-BFP-SKL-HIS3 PEX13::msGFP-HphNT1* | This study |
| ΔPex11 TEF1pr-Pex3 Erg6-2xmKate2 pRS403 TEFpr-BFP-SKL Pex14-msGFP | AGMY1779 | *BY4741 – Pex11Δ::KanMX PEX3pr::TEF1pr-NatNT2 ERG6::2xmKate2-URA3 HIS3::pRS403-TEF1pr-BFP-SKL-HIS3 PEX14::msGFP-HphNT1* | This study |
| TEF1pr-Pex3 Pex14-2xmKate2 | AGMY1258 | *BY4741 – PEX3pr::TEF1pr-NatNT2 PEX14::2xmKate2-HphNT1* | This study |
| TEF1pr-Pex3 Pex14-2xmKate2 | AGMY1303 | *W303 – PEX3pr::TEF1pr-NatNT2 PEX14::2xmKate2-URA3* | This study |
| ΔLDs | FFY30 | *W303 - Dga1::KanMX4 Liro1::TRP1 Are1::HIS3 Are2::LEU2 ADE2* | Froelich Lab |
| ΔLDs TEF1pr-Pex3 Pex14-2xmKate2 | AGMY1517 | *W303 - PEX3pr::TEF1pr-NatNT2 PEX14::2xmKate2-URA3* | This study |
| TEF1pr-Pex3 ΔPex35 Pex14-2xmKate2 | AGMY1466 | *BY4741 – PEX3pr::TEF1pr-NatNT2 Pex35Δ::HphNT1 PEX14::2xmKate2-URA3* | This study |
| TEF1pr-Pex3 ΔSap1 mCherry-SKL | AGMY1269 | *BY4741 – PEX3pr::TEF1pr-NatNT2 Sap1Δ::HphNT1 LEU2::mCherry-SKL-LEU2* | This study |
| TEF1pr-Pex3 ΔLdo45/16 Pex14-2xmKate2 | AGMY1467 | *BY4741 – PEX3pr::TEF1pr-NatNT2 Ldo45/16Δ::HphNT1 PEX14::2xmKate2-URA3* | This study |
| TEF1pr-Pex3 mCherry-SKL ΔPex30 | AGMY1929 | *BY4741 – PEX3pr::TEF1pr-NatNT2 LEU2::mCherry-SKL-LEU2 Pex30Δ::HphNT1* | This study |
| TEF1pr-Pex3 mCherry-SKL ΔPex34 | AGMY1933 | *BY4741 – PEX3pr::TEF1pr-NatNT2 LEU2::mCherry-SKL-LEU2 Pex34Δ::HphNT1* | This study |
| TEF1pr-Pex3 mCherry-SKL ΔFzo1 | AGMY1928 | *BY4741 – PEX3pr::TEF1pr-NatNT2 LEU2::mCherry-SKL-LEU2 Fzo1Δ::HphNT1* | This study |
| Erg6-2xmKate2 Pex14-HaloTag pRS403 TEF-GFP-Pex3(CD) | AGMY1615 | *BY4741 – ERG6::2xmKate2-URA3 PEX14::HaloTag-MET17 HIS3::pRS403-TEF1pr-GFP-PEX3(CD)-HIS3* | This study |
| Tom70-NbALFA Cit1-Halo GFP-SKL pRS403 TEF1pr-ALFATag-mKate2-Pex3(CD) | AGMY1851 | *BY4741 – TOM70::NbALFA-KanMX CIT1-HaloTag-MET17 LEU2::GFP-SKL-LEU2 HIS3:: pRS403-TEF1pr-ALFATag-mKate2-PEX3(CD)-HIS3* | This study |
| TEF1pr-Pex3 Pex14-2xmKate2 ΔInp1 | AGMY1536 | BY4741 – PEX3pr::TEF1pr-NatNT2 PEX14::2xmKate2-HphNT1 Inp1Δ::KanMX | This study |
| TEF1pr-Pex3 Pex14-2xmKate2 ΔAtg36 | AGMY1582 | BY4741 – PEX3pr::TEF1pr-NatNT2 PEX14::2xmKate2-HphNT1 Atg36Δ::KanMX | This study |
| ΔPex3 pRS403 TEF1pr-Pex3(WT)-mKate2 BFP-SKL | AGMY1896 | BY4741 – Pex3Δ::HphNT1 HIS3::pRS403-TEF1pr-PEX3(WT)-mKate2-HIS3 LEU2::BFP-SKL-LEU2 | This study |
| ΔPex3 pRS403 TEF1pr-Pex3(W128K, L131K)-mKate2 BFP-SKL | AGMY1897 | BY4741 – Pex3Δ::HphNT1 HIS3::pRS403-TEF1pr-PEX3(W128K, L131K)-mKate2-HIS3 LEU2::BFP-SKL-LEU2 | This study |
| TEF1pr-msGFP2-Pex19 pRS403 TEF1pr-Pex3(WT)-mKate2-ALFATag | AGMY1919 | BY4741 – PEX19pr::TEF1pr-msGFP2-NatNT1 HIS3::pRS403-TEF1pr-PEX3(WT)-mKate2-ALFATag-HIS3 | This study |
| TEF1pr-msGFP2-Pex19 pRS403 TEF1pr-Pex3(W128K, L131K)-mKate2-ALFATag | AGMY1920 | BY4741 – PEX19pr::TEF1pr-msGFP2-NatNT1 HIS3::pRS403-TEF1pr-PEX3(W128K, L131K)-mKate2-ALFATag-HIS3 | This study |
| pRS403 TEF1pr-Pex3(WT)-mKate2 | AGMY1820 | BY4741 – HIS3::pRS403-TEF1pr-PEX3(WT)-mKate2 | This study |
| pRS403 TEF1pr-Pex3(W128K, L131K)-mKate2 | AGMY1821 | BY4741 – HIS3::pRS403-TEF1pr-PEX3(W128K, L131K)-mKate2 | This study |
| TEF1pr-Pex3 mCherry-SKL ΔTgl4 | AGMY1729 | BY4741 – PEX3pr::TEF1pr-NatNT2 LEU2::mCherry-SKL-LEU2 Tgl4Δ::HphNT1 | This study |
| TEF1pr-Pex3 mCherry-SKL ΔTgl3 | AGMY1844 | BY4741 – PEX3pr::TEF1pr-NatNT2 LEU2::mCherry-SKL-LEU2 Tgl3Δ::KanMX | This study |
| TEF1pr-Pex3 mCherry-SKL ΔTgl5 | AGMY1810 | BY4741 – PEX3pr::TEF1pr-NatNT2 LEU2::mCherry-SKL-LEU2 Tgl5Δ::KanMX | This study |
| TEF1pr-Pex3 mCherry-SKL ΔTgl1 | AGMY1926 | BY4741 – PEX3pr::TEF1pr-NatNT2 LEU2::mCherry-SKL-LEU2 Tgl1Δ::HphNT1 | This study |
| TEF1pr-Pex3 mCherry-SKL ΔLdh1 | AGMY1927 | BY4741 – PEX3pr::TEF1pr-NatNT2 LEU2::mCherry-SKL-LEU2 Ldh1Δ::HphNT1 | This study |
| TEF1pr-Pex3 mCherry-SKL ΔYeh1 | AGMY1936 | BY4741 – PEX3pr::TEF1pr-NatNT2 LEU2::mCherry-SKL-LEU2 Yeh1Δ::HphNT1 | This study |
| ΔPex11 TEF1pr-Pex3 Erg6-2xmKate2 pRS403 TEF1pr-BFP-SKL Tgl4-msGFP | AGMY1782 | BY4741 – pex11Δ::KanMX PEX3pr::TEF1pr-NatNT2 ERG6::2xmKate2-URA3 HIS3::pRS403-TEF1pr-BFP-SKL-HIS3 TGL4::msGFP-HphNT1 | This study |
| ΔPex11 TEF1pr-Pex3 Erg6-2xmKate2 pRS403 TEF1pr-BFP-SKL Tgl3-msGFP | AGMY1823 | BY4741 – pex11Δ::KanMX PEX3pr::TEF1pr-NatNT2 ERG6::2xmKate2-URA3 HIS3::pRS403-TEF1pr-BFP-SKL-HIS3 TGL3::msGFP-HphNT1 | This study |
| ΔPex11 TEF1pr-Pex3 Erg6-2xmKate2 pRS403 TEF1pr-BFP-SKL Tgl5-msGFP | AGMY1780 | BY4741 – pex11Δ::KanMX PEX3pr::TEF1pr-NatNT2 ERG6::2xmKate2-URA3 HIS3::pRS403-TEF1pr-BFP-SKL-HIS3 TGL5::msGFP-HphNT1 | This study |
| Tgl4 (S315G) | AGMY1888 | BY4741 – TGL4 (S315G) | This study |
| Tgl4 (S315G) mCherry-SKL TEF1pr-Pex3 | AGMY1895 | BY4741 – TGL4 (S315G) LEU2::mCherry-SKL-LEU2 | This study |
| TEF2pr-Pex3 Erg6-mCherry BFP-SKL Tgl4-GFP | YMB1326 | his3Δ1, leu2Δ0, lys2+/lys+, met15Δ0, ura3Δ0::BFP-SKL HIS3, can1Δ::STE2pr-sp HIS5, lyp1Δ::STE3pr-LEU2, Nat pTEF2-PEX3, TGL4-GFP URA3, ERG6-mCherry HYG | This study |

**Supplemental Table 2:** Plasmids used in this study.

| Name | Identifier | Description | Yeast selection marker | Source |
| --- | --- | --- | --- | --- |
| pRS403 TEF1pr-BFP-SKL | pAGM197 | Plasmid to be integrated in the *HIS3* locus. It expresses BFP with a Pts1 signal to target it to the lumen of peroxisomes. | HIS3MX6 | This study |
| pRS403 TEF1pr-GFP-Pex3(CD) | pAGM134 | Plasmid to express Pex3 (CD) with an N-terminal GFP tag. | HIS3MX6 | This study |
| pRS403 TEF1pr-AlfaTag-mKate2-Pex3(CD) | pAGM212 | Plasmid to express Pex3 (CD) with an N-terminal ALFA and mKate2 tag. | HIS3MX6 | This study |
| pRS403 TEF1pr-Pex3-mKate2 | pAGM210 | Plasmid to express the full-length Pex3 under the control of the *TEF1* promoter, with a C-terminal mKate2 tag. | HIS3MX6 | This study |
| pRS403 TEF1pr-Pex3 (W128K, L131K)-mKate2 | pAGM211 | Plasmid to express the mutant variant of Pex3 (which does not interact with Pex19), under the control of the *TEF1* promoter, with a C-terminal mKate2 tag. | HIS3MX6 | This study |
| pRS403 TEF1pr-Pex3-mKate2-Alfatag | pAGM225 | Plasmid to express the full-length Pex3 under the control of the *TEF1* promoter, with a C-terminal mKate2 and ALFA tag. | HIS3MX6 | This study |
| pRS403 TEF1pr-Pex3 (W128K, L131K)-mKate2-Alfatag | pAGM226 | Plasmid to express Pex3(W128K, L131K) under the control of the *TEF1* promoter with a C-terminal tag consisting of mKate2 and ALFA tag. | HIS3MX6 | This study |
| pRS315 mCherry-SKL | CU5065 | Plasmid to express mCherry with a Pts1 signal to target it to the lumen of peroxisomes. | LEU2 | Gift from Judith Müller (Institute of Molecular Genetics and Cell Biology, Ulm University) |
| pRS415 GFP-SKL | CU5066 | Plasmid to express GFP with a Pts1 signal to target it to the lumen of peroxisomes. | LEU2 | (Schäfer et al., 2004) |
| pRS415 TEF1pr-BFP-SKL | pAGM224 | Plasmid to express BFP with a Pts1 signal to target it to the lumen of peroxisomes. | LEU2 | This study |
| pRS403 TGL4pr-Tgl4-msGFP2 | pAGM242 | Plasmid to express Tgl4 under its endogenous promoter, with a C-terminal msGFP2 tag. | HIS3MX6 | This study |

**Supplemental Table 3:**
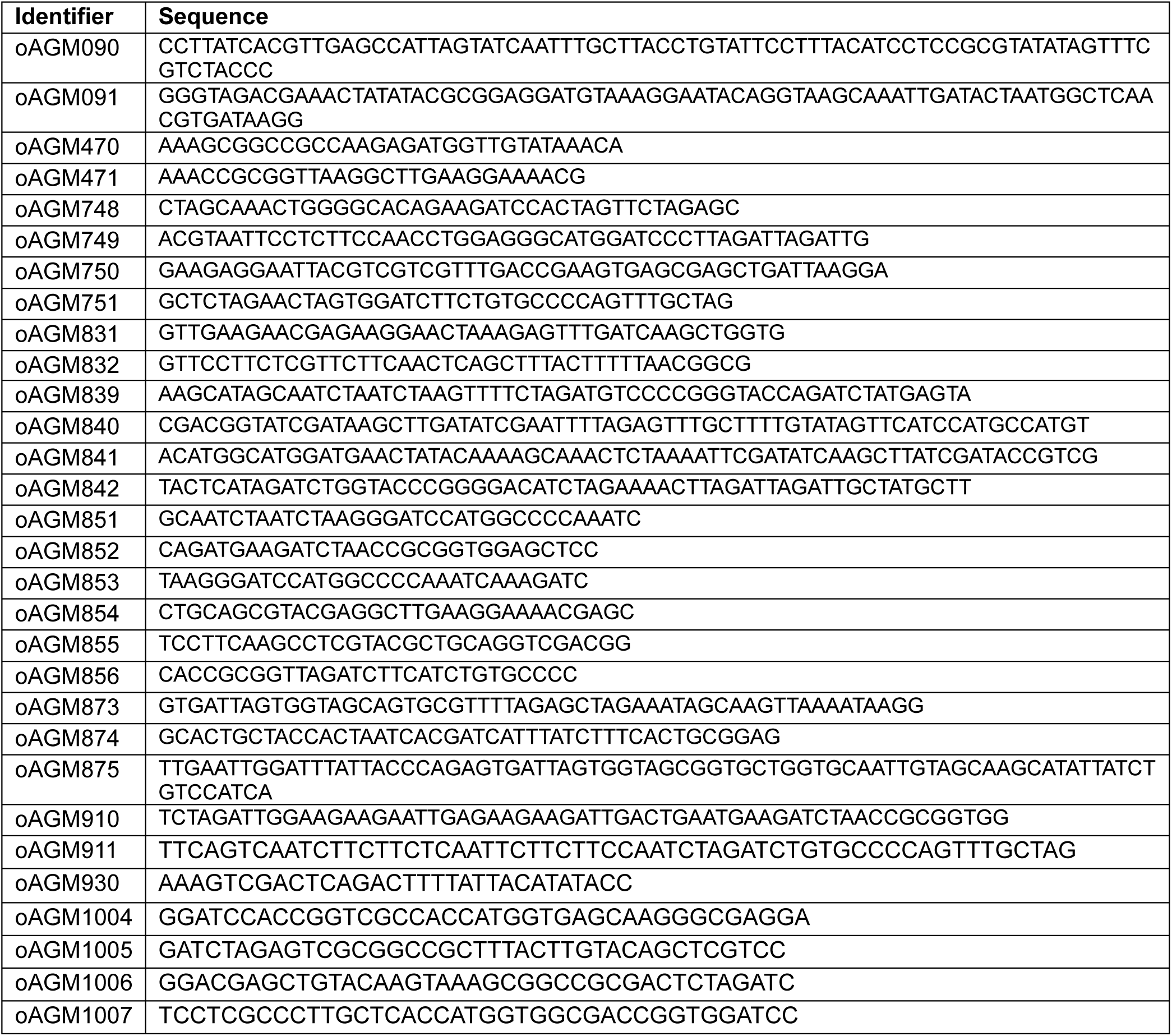
Oligonucleotides used in this study.

**Supplemental Table 4:**
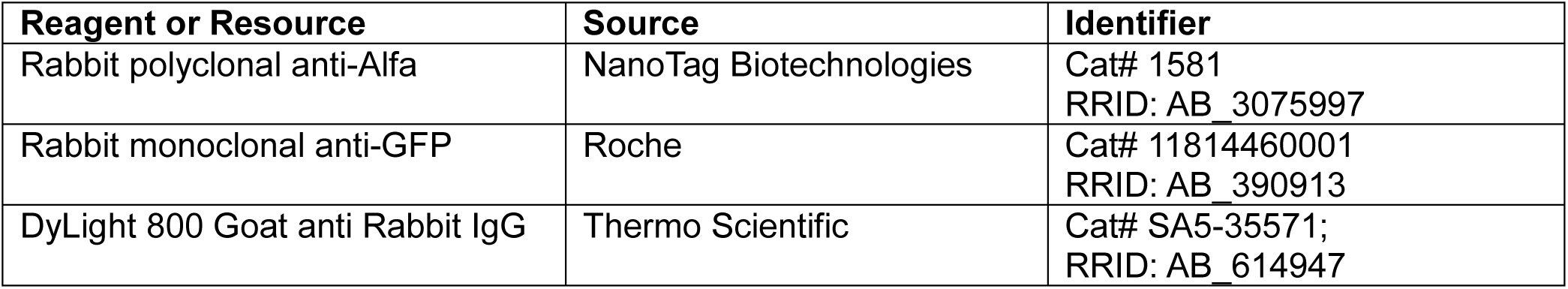
Antibodies used in this study.

| Reagent or Resource | Source | Identifier |
| --- | --- | --- |
| Rabbit polyclonal anti-Alfa | NanoTag Biotechnologies | Cat# 1581<br>RRID: AB_3075997 |
| Rabbit monoclonal anti-GFP | Roche | Cat# 11814460001<br>RRID: AB_390913 |
| DyLight 800 Goat anti Rabbit IgG | Thermo Scientific | Cat# SA5-35571;<br>RRID: AB_614947 |

**Supplemental Table 5:** Hits from genome-wide screen for deletions that disrupt Tgl4 localization at lipid droplet-peroxisome contact site.

| Systematic name | Standard name |
| --- | --- |
| YOR037W | CYC2 |
| YOR374W | ALD4 |
| YLR377C | FBP1 |
| YGL218W |  |
| YLR393W | ATP10 |
| YLR278C |  |
| YPL060W | MFM1 |
| YFR033C | QCR6 |
| YPR124W | CTR1 |
| YAL048C | GEM1 |
| YML060W | OGG1 |
| YML030W | RCF1 |
| YMR214W | SCJ1 |
| YMR243C | ZRC1 |
| YMR245W |  |
| YMR300C | ADE4 |
| YOL003C | PFA4 |
| YOR086C | TCB1 |
| YDR392W | SPT3 |
| YBR204C | LDH1 |
| YDR116C | MRPL1 |
| YBR238C |  |
| YGR183C | QCR9 |
| YHR011W | DIA4 |
| YKL106W | AAT1 |
| YLR218C | COA4 |
| YOR221C | MCT1 |
| YKL137W | CMC1 |
| YGR118W | RPS23A |
| YKL032C | IXR1 |
| YKL170W | MRPL38 |
| YGL246C | RAI1 |
| YDR207C | UME6 |
| YPL079W | RPL21B |
| YPR188C | MLC2 |
| YNL200C | NNR1 |
| YGL026C | TRP5 |
| YDR295C | HDA2 |
| YDR320C | SWA2 |
| YKR016W | MIC60 |
| YGR279C | SCW4 |
| YLR038C | COX12 |
| YJR120W | DMO1 |
| YML072C | TCB3 |
| YJR077C | MIR1 |
| YDL172C |  |
| YBR131W | CCZ1 |
| YJL128C | PBS2 |
| YGL127C | SOH1 |
| YER182W | FMP10 |
| YGL148W | ARO2 |
| YBR299W | MAL32 |
| YCR107W | AAD3 |
| YFL004W | VTC2 |
| YFL012W |  |
| YER163C | GCG1 |
| YGR273C |  |
| YER166W | DNF1 |
| YGR289C | MAL11 |
| YGR292W | MAL12 |
| YLR226W | BUR2 |
| YNL055C | POR1 |
| YPL050C | MNN9 |
| YBL002W | HTB2 |
| YGL070C | RPB9 |
| YNL069C | RPL16B |
| YJL003W | COX16 |
| YJL027C |  |
| YJL028W |  |
| YJR004C | SAG1 |
| YJR037W |  |
| YDL167C | NRP1 |
| YNL003C | PET8 |
| YNL084C | END3 |
| YER122C | GLO3 |
| YJL075C | APQ13 |
| YOL052C-A | DDR2 |
| YOL077W-A | ATP19 |
| YMR156C | TPP1 |
| YLR099C | ICT1 |

| **Systematic name** | **Standard name** |
| --- | --- |
| YOR087W | YVC1 |
| YDR389W | SAC7 |
| YDR422C | SIP1 |
| YDR351W | SBE2 |
| YMR063W | RIM9 |
| YMR251W-A | HOR7 |

## Notes

### Competing Interest Statement

The authors have declared no competing interest.

## REFERENCES

Ader, N. R., & Kukulski, W. (2017). triCLEM: Combining high-precision, room temperature CLEM with cryo-fluorescence microscopy to identify very rare events. Methods in Cell Biology, 140, 303–320. 10.1016/bs.mcb.2017.03.009

Álvarez-Guerra, I., Block, E., Broeskamp, F., Gabrijelčič, S., Infant, T., de Ory, A., Habernig, L., Andréasson, C., Levine, T. P., Höög, J. L., & Büttner, S. (2024). LDO proteins and Vac8 form a vacuole-lipid droplet contact site to enable starvation-induced lipophagy in yeast. Developmental Cell, 59(6), 759–775.e5. 10.1016/j.devcel.2024.01.014

Athenstaedt, K., & Daum, G. (2003). YMR313c/TGL3 encodes a novel triacylglycerol lipase located in lipid particles of Saccharomyces cerevisiae. Journal of Biological Chemistry, 278(26), 23317– 23323. 10.1074/jbc.M302577200

Athenstaedt, K., & Daum, G. (2005). Tgl4p and Tgl5p, two triacylglycerol lipases of the yeast Saccharomyces cerevisiae are localized to lipid particles. Journal of Biological Chemistry, 280(45), 37301–37309. 10.1074/jbc.M507261200

Binns, D., Januszewski, T., Chen, Y., Hill, J., Markin, V. S., Zhao, Y., Gilpin, C., Chapman, K. D., Anderson, R. G. W., & Goodman, J. M. (2006). An intimate collaboration between peroxisomes and lipid bodies. Journal of Cell Biology, 173(5), 719–731. 10.1083/jcb.200511125

Bonekamp, N. A., Sampaio, P., de Abreu, F. V., Lüers, G. H., & Schrader, M. (2012). Transient Complex Interactions of Mammalian Peroxisomes Without Exchange of Matrix or Membrane Marker Proteins. Traffic, 13(7), 960–978. 10.1111/j.1600-0854.2012.01356.x

Brand, H., & Perrimon, N. (1993). Targeted gene expression as a means of altering cell fates and generating dominant phenotypes.

Breslow, D. K., Cameron, D. M., Collins, S. R., Schuldiner, M., Stewart-Ornstein, J., Newman, H. W., Braun, S., Madhani, H. D., Krogan, N. J., & Weissman, J. S. (n.d.). A comprehensive strategy enabling high-resolution functional analysis of the yeast genome.

Burnett, S. F., Farré, J. C., Nazarko, T. Y., & Subramani, S. (2015). Peroxisomal Pex3 activates selective autophagy of peroxisomes via interaction with the pexophagy receptor Atg30. Journal of Biological Chemistry, 290(13), 8623–8631. 10.1074/jbc.M114.619338

Chang, C. L., Weigel, A. V., Ioannou, M. S., Amalia Pasolli, H., Shan Xu, C., Peale, D. R., Shtengel, G., Freeman, M., Hess, H. F., Blackstone, C., & Lippincott-Schwartz, J. (2019). Spastin tethers lipid droplets to peroxisomes and directs fatty acid trafficking through ESCRT-III. Journal of Cell Biology, 218(8), 2583–2599. 10.1083/jcb.201902061

Cohen, Y., & Schuldiner, M. (2011). Advanced Methods for High-Throughput Microscopy Screening of Genetically Modified Yeast Libraries. In Methods in Molecular Biology (Vol. 781, pp. 127–159). Humana Press Inc. 10.1007/978-1-61779-276-2_8

Cox, J., & Mann, M. (2008). MaxQuant enables high peptide identification rates, individualized p.p.b.-range mass accuracies and proteome-wide protein quantification. Nature Biotechnology, 26(12), 1367–1372. 10.1038/nbt.1511

Cox, J., Neuhauser, N., Michalski, A., Scheltema, R. A., Olsen, J. V., & Mann, M. (2011). Andromeda: A peptide search engine integrated into the MaxQuant environment. Journal of Proteome Research, 10(4), 1794–1805. 10.1021/pr101065j

David, C., Koch, J., Oeljeklaus, S., Laernsack, A., Melchior, S., Wiese, S., Schummer, A., Erdmann, R., Warscheid, B., & Brocard, C. (2013). A combined approach of quantitative interaction proteomics and live-cell imaging reveals a regulatory role for endoplasmic reticulum (ER) Reticulon Homology Proteins in peroxisome biogenesis. Molecular and Cellular Proteomics, 12(9), 2408–2425. 10.1074/mcp.M112.017830

Day, K. J., Casler, J. C., & Glick, B. S. (2018). Budding Yeast Has a Minimal Endomembrane System. Developmental Cell, 44(1), 56–72.e4. 10.1016/j.devcel.2017.12.014

Diep, D. T. V., Collado, J., Hugenroth, M., Fausten, R. M., Percifull, L., Wälte, M., Schuberth, C., Schmidt, O., Fernández-Busnadiego, R., & Bohnert, M. (2024). A metabolically controlled contact site between vacuoles and lipid droplets in yeast. Developmental Cell, 59(6), 740–758.e10. 10.1016/j.devcel.2024.01.016

Eisenberg-Bord, M., Shai, N., Schuldiner, M., & Bohnert, M. (2016). A Tether Is a Tether Is a Tether: Tethering at Membrane Contact Sites. Developmental Cell, 39(4), 395–409. 10.1016/j.devcel.2016.10.022

Elbaz-Alon, Y., Eisenberg-Bord, M., Shinder, V., Stiller, S. B., Shimoni, E., Wiedemann, N., Geiger, T., & Schuldiner, M. (2015). Lam6 Regulates the Extent of Contacts between Organelles. Cell Reports, 12(1), 7–14. 10.1016/j.celrep.2015.06.022

Erdmann, R., & Blobel, G. (1995). Giant peroxisomes in oleic acid-induced Saccharomyces cerevisiae lacking the peroxisomal membrane protein Pmp27p. Journal of Cell Biology, 128(4), 509–523. 10.1083/jcb.128.4.509

Fang, Y., Morrell, J. C., Jones, J. M., & Gould, S. J. (2004). PEX3 functions as a PEX19 docking factor in the import of class I peroxisomal membrane proteins. Journal of Cell Biology, 164(6), 863–875. 10.1083/jcb.200311131

Filadi, R., Leal, N. S., Schreiner, B., Rossi, A., Dentoni, G., Pinho, C. M., Wiehager, B., Cieri, D., Calì, T., Pizzo, P., & Ankarcrona, M. (2018). TOM70 Sustains Cell Bioenergetics by Promoting IP3R3-Mediated ER to Mitochondria Ca2+ Transfer. Current Biology, 28(3), 369–382.e6. 10.1016/j.cub.2017.12.047

Generoso, W. C., Gottardi, M., Oreb, M., & Boles, E. (2016). Simplified CRISPR-Cas genome editing for Saccharomyces cerevisiae. Journal of Microbiological Methods, 127, 203–205. 10.1016/j.mimet.2016.06.020

Giaever, G., Chu, A. M., Ni, L., Connelly, C., Riles, L., Vé ronneau, S., Dow, S., Lucau-Danila, A., Anderson, K., André, B., Arkin, A. P., Astromoff, A., El Bakkoury, M., Bangham, R., Benito, R., Brachat, S., Campanaro, S., Curtiss, M., Davis, K., … Johnston, M. (2002). Functional profiling of the Saccharomyces cerevisiae genome. http://www.kegg.com

González Montoro, A., Vargas Duarte, P., Auffarth, K., Walter, S., Fröhlich, F., & Ungermann, C. (2021). Subunit exchange among endolysosomal tethering complexes is linked to contact site formation at the vacuole. Molecular Biology of the Cell, 32(22), br14. 10.1091/mbc.E21-05-0227

Gorgas, K., & Zaar, K. (1984). Anatomy and Embryology Peroxisomes in sebaceous glands III. Morphological similarities of peroxisomes with smooth endoplasmic reticulum and Golgi stacks in the circumanal gland of the dog* **. In Anat Embryol (Vol. 169).

Götzke, H., Kilisch, M., Martínez-Carranza, M., Sograte-Idrissi, S., Rajavel, A., Schlichthaerle, T., Engels, N., Jungmann, R., Stenmark, P., Opazo, F., & Frey, S. (2019). The ALFA-tag is a highly versatile tool for nanobody-based bioscience applications. Nature Communications, 10(1), 1–12. 10.1038/s41467-019-12301-7

Grimm, J. B., Xie, L., Casler, J. C., Patel, R., Tkachuk, A. N., Falco, N., Choi, H., Lippincott-Schwartz, J., Brown, T. A., Glick, B. S., Liu, Z., & Lavis, L. D. (2021). A General Method to Improve Fluorophores Using Deuterated Auxochromes. JACS Au, 1(5), 690–696. 10.1021/jacsau.1c00006

Hettema, E. H., Girzalsky, W., Van Den Berg, M., Erdmann, R., & Distel, B. (2000). Saccharomyces cerevisiae Pex3p and Pex19p are required for proper localization and stability of peroxisomal membrane proteins. EMBO Journal, 19(2), 223–233. 10.1093/emboj/19.2.223

Hin, A., Tong, Y., & Boone, C. (2006). Synthetic Genetic Array Analysis in Saccharomyces cerevisiae. In From: Methods in Molecular Biology (Vol. 313).

Höhfeld, J., Veenhuis, M., & Kunau, W. H. (1991). PAS3, a Saccharomyces cerevisiae gene encoding a peroxisomal integral membrane protein essential for peroxisome biogenesis. Journal of Cell Biology, 114(6), 1167–1178. 10.1083/jcb.114.6.1167

Hollenstein, D. M., Gómez-Sánchez, R., Ciftci, A., Kriegenburg, F., Mari, M., Torggler, R., Licheva, M., Reggiori, F., & Kraft, C. (2019). Vac8 spatially confines autophagosome formation at the vacuole in S. Cerevisiae. Journal of Cell Science, 132(22). 10.1242/jcs.235002

Hulmes, G. E., Hutchinson, J. D., Dahan, N., Nuttall, J. M., Allwood, E. G., Ayscough, K. R., & Hettema, E. H. (2020). The Pex3-Inp1 complex tethers yeast peroxisomes to the plasma membrane. Journal of Cell Biology, 219(10). 10.1083/JCB.201906021

Jandrositz, A., Petschnigg, J., Zimmermann, R., Natter, K., Scholze, H., Hermetter, A., Kohlwein, S. D., & Leber, R. (2005). The lipid droplet enzyme Tgl1p hydrolyzes both steryl esters and triglycerides in the yeast, Saccharomyces cerevisiae. Biochimica et Biophysica Acta - Molecular and Cell Biology of Lipids, 1735(1), 50–58. 10.1016/j.bbalip.2005.04.005

Janke, C., Magiera, M. M., Rathfelder, N., Taxis, C., Reber, S., Maekawa, H., Moreno-Borchart, A., Doenges, G., Schwob, E., Schiebel, E., & Knop, M. (2004). A versatile toolbox for PCR-based tagging of yeast genes: New fluorescent proteins, more markers and promoter substitution cassettes. Yeast, 21(11), 947–962. 10.1002/yea.1142

Kakimoto, Y., Tashiro, S., Kojima, R., Morozumi, Y., Endo, T., & Tamura, Y. (2018). Visualizing multiple inter-organelle contact sites using the organelle-Targeted split-GFP system. Scientific Reports, 8(1). 10.1038/s41598-018-24466-0

Kammerer, S., Holzinger, A., Welsch, U., & Roscher, A. A. (1998). Cloning and characterization of the gene encoding the human peroxisomal assembly protein Pex3p. FEBS Letters, 429(1), 53–60. 10.1016/S0014-5793(98)00557-2

Knoblach, B., Sun, X., Coquelle, N., Fagarasanu, A., Poirier, R. L., & Rachubinski, R. A. (2013a). An ER-peroxisome tether exerts peroxisome population control in yeast. EMBO Journal, 32(18), 2439–2453. 10.1038/emboj.2013.170

Knoblach, B., Sun, X., Coquelle, N., Fagarasanu, A., Poirier, R. L., & Rachubinski, R. A. (2013b). An ER-peroxisome tether exerts peroxisome population control in yeast. EMBO Journal, 32(18), 2439–2453. 10.1038/emboj.2013.170

Köffel, R., Tiwari, R., Falquet, L., & Schneiter, R. (2005). The Saccharomyces cerevisiae YLL012/YEH1, YLR020/YEH2, and TGL1 Genes Encode a Novel Family of Membrane-Anchored Lipases That Are Required for Steryl Ester Hydrolysis. Molecular and Cell Biology, 25(5), 1655–1668. 10.1128/MCB.25.5.1655

Kurat, C. F., Natter, K., Petschnigg, J., Wolinski, H., Scheuringer, K., Scholz, H., Zimmermann, R., Leber, R., Zechner, R., & Kohlwein, S. D. (2006). Obese yeast: Triglyceride lipolysis is functionally conserved from mammals to yeast. Journal of Biological Chemistry, 281(1), 491–500. 10.1074/jbc.M508414200

Kvam, E., & Goldfarb, D. S. (2004). Nvj1p is the outer-nuclear-membrane receptor for oxysterol-binding protein homolog Osh1p in Saccharomyces cerevisiae. Journal of Cell Science, 117(21), 4959– 4968. 10.1242/jcs.01372

Levine, T. P., & Munro, S. (2001). Dual targeting of Osh1p, a yeast homologue of oxysterol-binding protein, to both the Golgi and the nucleus-vacuole junction. Molecular Biology of the Cell, 12(6), 1633–1644. 10.1091/mbc.12.6.1633

Liu, L. K., Choudhary, V., Toulmay, A., & Prinz, W. A. (2017). An inducible ER-Golgi tether facilitates ceramide transport to alleviate lipotoxicity. Journal of Cell Biology, 216(1), 131–147. 10.1083/jcb.201606059

Loewen, C. J. R., Roy, A., & Levine, T. P. (2003). A conserved ER targeting motif in three families of lipid binding proteins and in Opi1p binds VAP. EMBO Journal, 22(9), 2025–2035. 10.1093/emboj/cdg201

Marta Fernández-Suárez, T. Scott Chen, and A. Y. T. (2008). Protein-Protein Interaction Detection In Vitro and in Cells by Proximity Biotinylation. NIH Public Access, 23(1), 1–7. 10.1021/ja801445p.Protein-Protein

Mast, F. D., Rachubinski, R. A., & Aitchison, J. D. (2020). Peroxisome prognostications: Exploring the birth, life, and death of an organelle. Journal of Cell Biology, 219(3), 1–13. 10.1083/JCB.201912100

Motley, A. M., Nuttall, J. M., & Hettema, E. H. (2012). Pex3-anchored Atg36 tags peroxisomes for degradation in Saccharomyces cerevisiae. EMBO Journal, 31(13), 2852–2868. 10.1038/emboj.2012.151

Murley, A., Sarsam, R. D., Toulmay, A., Yamada, J., Prinz, W. A., & Nunnari, J. (2015). Ltc1 is an ER-localized sterol transporter and a component of ER-mitochondria and ER-vacuole contacts. Journal of Cell Biology, 209(4), 539–548. 10.1083/jcb.201502033

Ollion, J., Cochennec, J., Loll, F., Escudé, C., & Boudier, T. (2013). TANGO: A generic tool for high-throughput 3D image analysis for studying nuclear organization. Bioinformatics, 29(14), 1840– 1841. 10.1093/bioinformatics/btt276

Olsen, J. V., Macek, B., Lange, O., Makarov, A., Horning, S., & Mann, M. (2007). Higher-energy C-trap dissociation for peptide modification analysis. Nature Methods, 4(9), 709–712. 10.1038/nmeth1060

Pan, X., Roberts, P., Chen, Y., Kvam, E., Shulga, N., Huang, K., Lemmon, S., & Goldfarb, D. S. (2000). Nucleus-vacuole junctions in Saccharomyces cerevisiae are formed through the direct interaction of Vac8p with Nvj1p. Molecular Biology of the Cell, 11(7), 2445–2457. 10.1091/mbc.11.7.2445

Prinz, W. A., Toulmay, A., & Balla, T. (2020). The functional universe of membrane contact sites. Nature Reviews Molecular Cell Biology, 21(1), 7–24. 10.1038/s41580-019-0180-9

Rajakumari, S., Rajasekharan, R., & Daum, G. (2010). Triacylglycerol lipolysis is linked to sphingolipid and phospholipid metabolism of the yeast Saccharomyces cerevisiae. Biochimica et Biophysica Acta - Molecular and Cell Biology of Lipids, 1801(12), 1314–1322. 10.1016/j.bbalip.2010.08.004

S. Bolte & F. P. Cordelieres. (2006). A guided tour into subcellular colocalization analysis in light microscopy. In Journal of Microscopy (Vol. 224).

Sandager, L., Gustavsson, M. H., Ståhl, U., Dahlqvist, A., Wiberg, E., Banas, A., Lenman, M., Ronne, H., & Stymne, S. (2002). Storage lipid synthesis is non-essential in yeast. Journal of Biological Chemistry, 277(8), 6478–6482. 10.1074/jbc.M109109200

Sato, Y., Shibata, H., Nakano, H., Matsuzono, Y., Kashiwayama, Y., Kobayashi, Y., Fujiki, Y., Imanaka, T., & Kato, H. (2008). Characterization of the interaction between recombinant human peroxin Pex3p and Pex19p: Identification of TRP-104 in Pex3p as a critical residue for the interaction. Journal of Biological Chemistry, 283(10), 6136–6144. 10.1074/jbc.M706139200

Sato, Y., Shibata, H., Nakatsu, T., Nakano, H., Kashiwayama, Y., Imanaka, T., & Kato, H. (2010). Structural basis for docking of peroxisomal membrane protein carrier Pex19p onto its receptor Pex3p. EMBO Journal, 29(24), 4083–4093. 10.1038/emboj.2010.293

Schäfer, A., Kerssen, D., Veenhuis, M., Kunau, W.-H., & Schliebs, W. (2004). Functional Similarity between the Peroxisomal PTS2 Receptor Binding Protein Pex18p and the N-Terminal Half of the PTS1 Receptor Pex5p. Molecular and Cellular Biology, 24(20), 8895–8906. 10.1128/mcb.24.20.8895-8906.2004

Schmidt, F., Dietrich, D., Eylenstein, R., Groemping, Y., Stehle, T., & Dodt, G. (2012). The Role of Conserved PEX3 Regions in PEX19-Binding and Peroxisome Biogenesis. Traffic, 13(9), 1244– 1260. 10.1111/j.1600-0854.2012.01380.x

Schmidt, F., Treiber, N., Zocher, G., Bjelic, S., Steinmetz, M. O., Kalbacher, H., Stehle, T., & Dodt, G. (2010). Insights into peroxisome function from the structure of PEX3 in complex with a soluble fragment of PEX19. Journal of Biological Chemistry, 285(33), 25410–25417. 10.1074/jbc.M110.138503

Schmidt, O., Pfanner, N., & Meisinger, C. (2010). Mitochondrial protein import: From proteomics to functional mechanisms. In Nature Reviews Molecular Cell Biology (Vol. 11, Issue 9, pp. 655–667). 10.1038/nrm2959

Schrader, M., King, S. J., Stroh, T. A., & Schroer, T. A. (2000). Real time imaging reveals a peroxisomal reticulum in living cells. 3671, 3663–3671.

Scorrano, L., De Matteis, M. A., Emr, S., Giordano, F., Hajnóczky, G., Kornmann, B., Lackner, L. L., Levine, T. P., Pellegrini, L., Reinisch, K., Rizzuto, R., Simmen, T., Stenmark, H., Ungermann, C., & Schuldiner, M. (2019). Coming together to define membrane contact sites. Nature Communications, 10(1), 1–11. 10.1038/s41467-019-09253-3

Shai, N., Schuldiner, M., & Zalckvar, E. (2016). No peroxisome is an island - Peroxisome contact sites. Biochimica et Biophysica Acta - Molecular Cell Research, 1863(5), 1061–1069. 10.1016/j.bbamcr.2015.09.016

Shai, N., Yifrach, E., Van Roermund, C. W. T., Cohen, N., Bibi, C., Ijlst, L., Cavellini, L., Meurisse, J., Schuster, R., Zada, L., Mari, M. C., Reggiori, F. M., Hughes, A. L., Escobar-Henriques, M., Cohen, M. M., Waterham, H. R., Wanders, R. J. A., Schuldiner, M., & Zalckvar, E. (2018). Systematic mapping of contact sites reveals tethers and a function for the peroxisome-mitochondria contact. Nature Communications, 9(1). 10.1038/s41467-018-03957-8

Tess C. Branon, Justin A. Bosch, Ariana D. Sanchez, Namrata D. Udeshi, Tanya Svinkina, Steven A. Carr, Jessica L. Feldman, N. P. and A. Y. T. (2018). Efficient proximity labeling in living cells and organisms with TurboID. Nat Biotechnol., 176(3), 139–148. 10.1038/nbt.4201.Efficient

Thoms, S., Debelyy, M. O., Connerth, M., Daum, G., & Erdmann, R. (2011). The putative Saccharomyces cerevisiae hydrolase Ldh1p is localized to lipid droplets. Eukaryotic Cell, 10(6), 770–775. 10.1128/EC.05038-11

Toulmay, A., & Prinz, W. A. (2012). A conserved membrane-binding domain targets proteins to organelle contact sites. Journal of Cell Science, 125(1), 49–58. 10.1242/jcs.085118

Tyanova, S., Temu, T., & Cox, J. (2016). The MaxQuant computational platform for mass spectrometry-based shotgun proteomics. Nature Protocols, 11(12), 2301–2319. 10.1038/nprot.2016.136

Valm, A. M., Cohen, S., Legant, W. R., Melunis, J., Hershberg, U., Wait, E., Cohen, A. R., Davidson, M. W., Betzig, E., & Lippincott-Schwartz, J. (2017). Applying systems-level spectral imaging and analysis to reveal the organelle interactome. Nature, 546(7656), 162–167. 10.1038/nature22369

Veit, M., Laage, R., Dietrich, L., Wang, L., & Ungermann, C. (2001). Vac8p release from the SNARE complex and its palmitoylation are coupled and essential for vacuole fusion. EMBO Journal, 20(12), 3145–3155. 10.1093/emboj/20.12.3145

Voeltz, G. K., Sawyer, E. M., Hajnóczky, G., & Prinz, W. A. (2024). Making the connection: How membrane contact sites have changed our view of organelle biology. Cell, 187(2), 257–270. 10.1016/j.cell.2023.11.040

Wanders, R. J. A., Baes, M., Ribeiro, D., Ferdinandusse, S., & Waterham, H. R. (2023). the Physiological Functions of Human Peroxisomes. Physiological Reviews, 103(1), 957–1024. 10.1152/physrev.00051.2021

Wang, Y. X., Catlett, N. L., & Weisman, L. S. (1998). Vac8p, a vacuolar protein with armadillo repeats, functions in both vacuole inheritance and protein targeting from the cytoplasm to vacuole. Journal of Cell Biology, 140(5), 1063–1074. 10.1083/jcb.140.5.1063

Wu, H., de Boer, R., Krikken, A. M., Akşit, A., Yuan, W., & van der Klei, I. J. (2019a). Peroxisome development in yeast is associated with the formation of Pex3-dependent peroxisome-vacuole contact sites. Biochimica et Biophysica Acta - Molecular Cell Research, 1866(3), 349–359. 10.1016/j.bbamcr.2018.08.021

Wu, H., de Boer, R., Krikken, A. M., Akşit, A., Yuan, W., & van der Klei, I. J. (2019b). Peroxisome development in yeast is associated with the formation of Pex3-dependent peroxisome-vacuole contact sites. Biochimica et Biophysica Acta - Molecular Cell Research, 1866(3), 349–359. 10.1016/j.bbamcr.2018.08.021

Yamashita, S. ichi, Abe, K., Tatemichi, Y., & Fujiki, Y. (2014). The membrane peroxin PEX3 induces peroxisome-ubiquitination-linked pexophagy. Autophagy, 10(9), 1549–1564. 10.4161/auto.29329

Yan, M., Rachubinski, D. A., Joshi, S., Rachubinski, R. A., & Subramani, S. (2008). Dysferlin Domain-containing Proteins, Pex30p and Pex31p, Localized to Two Compartments, Control the Number and Size of Oleate-induced Peroxisomes in Pichia pastoris. Molecular Biology of the Cell, 19, 885–898. 10.1091/mbc.E07-10

Yifrach, E., Holbrook-Smith, D., Bürgi, J., Othman, A., Eisenstein, M., van Roermund, C. W., Visser, W., Tirosh, A., Rudowitz, M., Bibi, C., Galor, S., Weill, U., Fadel, A., Peleg, Y., Erdmann, R., Waterham, H. R., Wanders, R. J. A., Wilmanns, M., Zamboni, N., … Zalckvar, E. (2022). Systematic multi-level analysis of an organelle proteome reveals new peroxisomal functions. Molecular Systems Biology, 18(9), 1–21. 10.15252/msb.202211186

